# A novel taurine-respiring murine gut bacterium contributes to colonization resistance against enteropathogens

**DOI:** 10.1101/2022.10.05.510937

**Authors:** Huimin Ye, Sabrina Borusak, Claudia Eberl, Buck T. Hanson, Benjamin Zwirzitz, Craig W. Herbold, Petra Pjevac, Bela Hausmann, Bärbel Stecher, David Schleheck, Alexander Loy

## Abstract

Taurine-respiring gut bacteria produce H_2_S with ambivalent impact on host health. We report the isolation and genomic-ecophysiological characterization of the first taurine-respiring mouse gut bacterium. *Taurinivorans muris* represents a new widespread species with protective capacity against pathogens and differs from the human gut sulfidogen *Bilophila wadsworthia* in its sulfur metabolism and host distribution. Despite alternative physiologies, taurine respiration was the main *in vivo* lifestyle of *T. muris* independent of mouse diet and genotype. In gnotobiotic mice, *T. muris* selectively enhanced the activity of a sulfur metabolism gene-encoding prophage and provided slightly increased colonization resistance against *Salmonella* Typhimurium, which showed reduced expression of galactonate catabolism genes. We identified *T. muris* as the dominant sulfidogen of a mouse microbiota that conferred H_2_S-mediated protection against *Klebsiella pneumoniae* in a previous study. Together, we revealed the realized physiological niche of a key murine gut sulfidogen and its impact on pathogen and phage gene expression.

**One sentence summary:** Our work identified and characterized a new core member of the murine gut microbiota, revealed sulfidogenic taurine respiration as its predominant *in vivo* lifestyle, and emphasizes its protective function in pathogen colonization.

## Introduction

Hydrogen sulfide (H_2_S) is an intestinal metabolite with pleiotropic effects, particularly on the gut mucosa ^1,2^. H_2_S can have a detrimental impact on the intestinal epithelium by chemically disrupting the mucus barrier ^3^, causing DNA damage ^4^, and impairing energy generation in colonocytes through inhibition of cytochrome *c* oxidase and beta-oxidation of short-chain fatty acids ^5,6^. In contrast, low micromolar concentrations of H_2_S are anti-inflammatory and contribute to mucosal homeostasis and repair ^7,8^. Furthermore, H_2_S acts as a gaseous transmitter, a mitochondrial energy source, and an antioxidant in cellular redox processes, and thus its impact on mammalian physiology and health reaches beyond the gastrointestinal tract ^9^. For example, colonic luminal H_2_S can promote somatic pain in mice ^10^ and contribute to regulating blood pressure ^11,12^. The multiple (patho)physiological functions of H_2_S in various organs and tissues are dependent on its concentration and the health status of the host, but possibly also on the source of H_2_S. Mammalian cells can produce H_2_S from cysteine via several known pathways ^13^. In contrast to these endogenous sources, H_2_S-releasing drugs and H_2_S-producing intestinal microorganisms are considered exogenous sources. Compared to the colonic epithelium, sulfidogenic bacteria, which either metabolize organic sulfur compounds (e.g. cysteine) or anaerobically respire organic (e.g. taurine) or inorganic (e.g. sulfate, sulfite, tetrathionate) sulfur compounds in the gut, have a higher H_2_S-producing capacity and are thus potentially harmful to their hosts ^1,2,14^. Indeed, the abundance and activity of sulfidogenic gut bacteria were associated with intestinal diseases such as inflammatory bowel disease and colon cancer in various studies ^15–17^ and many gut pathogens, such as *Salmonella enterica* and *Chlostridioides difficile*, are also sulfidogenic ^18,19^. Excess bacterial H_2_S production combined with a reduced capacity of the inflamed mucosa to metabolize H_2_S is one of many mechanisms by which the gut microbiome can contribute to disease ^20^. Yet, the manifold endogenous and microbial factors and processes that regulate intestinal H_2_S homeostasis are insufficiently understood.

A major substrate of sulfidogenic bacteria in the gut is the organosulfonate taurine (2-aminoethanesulfonate) that derives directly from meat- or seafood-rich diets and is also liberated by microbial bile salt hydrolases (BSHs) from endogenously produced taurocholic bile acids ^21^. *Bilophila wadsworthia* is the most prominent taurine-utilizing bacterium in the human gut. Diets that contain high quantities of meat, dairy products or fats can be associated with the outgrowth of *B. wadsworthia* in the gut ^22,23^. Consumption of high-fat food triggers taurocholic bile acid production and increases the taurine: glycine ratio in the bile acid pool ^23^. In mouse models, higher abundances of *B. wadsworthia* can promote colitis and systemic inflammation ^23,24^ and aggravate metabolic dysfunctions ^25^.

*B. wadsworthia* metabolizes taurine via the two intermediates sulfoacetaldehyde and isethionate (2-hydroxyethanesulfonate) to sulfite ^26^, and the sulfite is utilized as electron acceptor for energy conservation and reduced to H_2_S via the DsrAB-DsrC dissimilatory sulfite reductase system ^27^. The highly oxygen-sensitive isethionate sulfite-lyase system IslAB catalyzes the abstraction of sulfite (desulfonation) of isethionate ^26,28^. Alternative taurine degradation pathways in other bacteria involve direct desulfonation of sulfoacetaldehyde by the oxygen-insensitive, thiamine-diphosphate-dependent sulfoacetaldehyde acetyltransferase Xsc ^29–31^. Xsc is employed in taurine utilization as carbon and energy source in a wide range of aerobic bacteria ^30,32^, as well as for anaerobic taurine fermentation by *Desulfonispora thiosulfatigenes* ^*31*^.

Here, we isolated the first taurine-respiring and H_2_S-producing bacterium from the murine intestinal tract and elucidated its fundamental and *in vivo* realized nutrient niche. Strain LT0009 represents a new *Desulfovibrionaceae* genus, termed *Taurinivorans muris* gen. nov., sp. nov., and differs from its human counterpart *B. wadsworthia* by using the Xsc pathway for taurine degradation and its distribution across different animal hosts. We further provide insights into the protective role of this newly identified species of the core mouse microbiome against pathogen colonization.

## Materials and methods

Supplementary Information provides further details on the methods described below.

### Isolation of strain LT0009 and growth experiments

Intestinal content of wild-type C57BL/6 mice was used as inoculum for the enrichment cultures. A modified *Desulfovibrio* medium was used for the isolation of strain LT0009 with taurine as electron acceptor and lactate and pyruvate as electron donors and for further growth experiments. Consumption of taurine and lactate and production of H_2_S and short-chain fatty acids (SCFA) were measured as previously reported ^33^.

### Microscopy

Gram staining of the LT0009 isolate was performed using a Gram-staining kit according to the manufacturer’s instruction (Sigma Aldrich, 77730-1KT-F) and its cellular morphology was imaged with a scanning electron microscope (JSM-IT300, JOEL). A new probe was designed, tested, and applied for fluorescence *in situ* hybridization (FISH)-based microscopy of the genus *Taurinivorans* (Supplementary Table 1, Supplementary Fig. 1).

### Genome sequencing and comparative sequence analyses

The complete genome of strain LT0009 was determined by combined short- (Illumina) and long-read (Nanopore) sequencing. The automated annotation of the genome was manually curated for genes of interest, focusing on energy metabolism. Phylogenomic analyses comprised treeing with 43 concatenated marker protein sequences and calculation of average amino acid identities (AAI) and whole-genome average nucleotide identities (gANI). Additional phylogenetic trees were calculated with LT0009 using sequences of the 16S rRNA gene and selected sulfur metabolism proteins or genes. Source information of 16S rRNA gene reference sequences was manually compiled from the NCBI SRA entries (Supplementary Table 2).

### Differential proteomics and transcriptomics

The total proteomes and transcriptomes of strain LT0009 grown with taurine, sulfolactate or thiosulfate as electron acceptor and lactate/pyruvate as electron donor were determined and compared.

### Analyses of publicly available 16S rRNA sequence data

The occurrence and prevalence of *Taurinivorans muris*- and *Bilophila wadsworthia*-related 16S rRNA gene sequences were analyzed across 123,723 amplicon datasets from gut samples, including 81,501 with host information. Further information on mouse studies with at least 20 samples that were positive for *B. wadsworthia* was manually compiled from the NCBI SRA entries or the corresponding publications (Supplementary Table 3).

The identity of *Desulfovibrionaceae* 16S rRNA gene sequences from amplicon sequencing data of wildR mice, which showed increased representation of *Deltaproteobacteria* and colonization resistance against the enteropathogen *Klebsiella pneumoniae* in a previous study ^34^, was re-analyzed.

### Gnotobiotic mouse experiments

The animal experiment was approved by the local authorities (Regierung von Oberbayern; ROB-55.2-2532.Vet_02-20-84). Twelve germ-free C57BL/6 mice were stably colonized with the 12-strain Oligo-Mouse-Microbiota (OMM^12^) community ^35^. OMM^12^ mice (n=6) were orally (50 µl) and rectally (100 µl) inoculated with the LT0009 strain. The control OMM^12^ mouse group (n=6) was treated with the same volume of sterile 1x phosphate-buffered saline. After 10 days, the mice were infected with the human enteric pathogen *Salmonella enterica* serovar Typhimurium (avirulent *S. enterica* Tm strain M2702; 5×10^7^c.f.u.) and sacrificed two days post infection (p. i.). Fecal microbiota composition was determined by strain-specific qPCR assays as previously described ^35^, including a newly developed assay for strain LT0009. Abundance of viable *S. enterica* Tm in feces and cecal content was determined by plating. Fecal samples of three mice from each group on day two p.i. were selected for metatranscriptome sequencing (JMF project JMF-2104-01) and analyses.

### LT0009-centric gut metatranscriptome analyses of laboratory mice

Cecal and fecal metatranscriptomes from a high-glucose diet experiment in mice (HG study, Hanson et al., unpublished) (JMP project JMF-2101-5) were analyzed for LT0009 gene expression. Mouse experiments were conducted following protocols approved by Austrian law (BMWF-66.006/ 0032-WF/V/3b/2014). Additionally, mouse gut metatranscriptomes from a previous study were analyzed for LT0009 gene expression (Plin2 study) ^36^. Sequence data (PRJNA379425) derived from eight-week-old C57BL/6 wild-type and Perilipin2-null (Plin2) mice fed high-fat/low-carbohydrate or low-fat/high-carbohydrate diets.

### Strain and data availability

Strain LT0009 has been deposited in the German Collection of Microorganisms and Cell Cultures (DSMZ) as DSM 111569 and the Japan Collection of Microorganisms (JCM) as JCM 34262. The genome and the 16S rRNA gene sequence of *T. muris* LT0009 are available at NCBI GenBank under accession numbers CP065938 and MW258658, respectively.

Sequencing data of the LT0009 pure culture transcriptome (JMF-2012-1) and the mouse gut metatranscriptomes from the HG study (JMF-2101-05) and the gnotobiotic study (JMF-2104-01) were deposited to the Sequence Read Archive (SRA) under BioProject accession PRJNA867178.

## Results and Discussion

### The first taurine-respiring bacterium isolated from the murine gut represents a new genus of the family *Desulfovibrionaceae*

Strain LT0009 was enriched from mouse gut contents (cecum and colon) using an anoxic, non-reducing, modified *Desulfovibrio* medium with L-lactate and pyruvate as electron donors (and carbon source) and taurine as the sulfite donor for sulfite respiration. Its isolation was achieved by several transfers in liquid medium, purification by streaking on ferric-iron supplemented agar plates, indicating sulfide production by black FeS formation and picking of black colonies, and by additional purification using dilution-to-extinction in liquid medium. Strain LT0009 produced sulfide and acetate during taurine degradation. We sequenced and reconstructed the complete LT0009 genome, which has a size of 2.2 Mbp, a G+C content of 43.6%, and is free of contamination as assessed by CheckM. The genome comprises 2,059 protein-coding genes, 56 tRNA genes, 4 rRNA operons (with 5S, 16S, and 23S rRNA genes), 4 pseudogenes, and 6 miscellaneous RNA genes.

LT0009 formed a monophyletic, genus-level (>94.5% similarity) lineage with other 16S rRNA gene sequences from the gut of mice and other hosts. It has <92% 16S rRNA gene sequence identity to the closest related isolates Marseille-P3669 and *Mailhella massiliensis* Marseille-P3199^T^, two strains isolated from human stool (Fig. 1a). Phylogenomic treeing and an AAI of <60% to other described species strongly suggested that LT0009 represents the type strain of a novel genus in the family *Desulfovibrionaceae* of the phylum *Desulfobacterota* ^*37*^ for which we propose the name *Taurinivorans muris* (Fig. 1b, Supplementary Fig. 2, Supplementary Information). The previously described mouse gut MAGs UBA8003 and extra-SRR4116659.59 have >98% ANI and AAI to LT0009 and thus would also belong to the species *T. muris* ^*38–40*^. Furthermore, the mouse gut MAGs extra-SRR4116662.45 and single-China-D-Fe10-120416.2 showed <80% AAI/ANI to strain LT0009 and 84% AAI and 82% ANI to each other, which indicates that each of these two MAGs would likely represent a separate species in the novel genus *Taurinivorans*. Notably, the genome of *T. muris* LT0009 with only 2.2 Mbp in size is considerably smaller than that of other free-living bacteria of the *Desulfovibrio-Bilophila-Lawsonia-Mailhella*-lineage (Fig. 1b). Only the obligate intracellular intestinal pathogen *Lawsonia intracellularis* has a smaller genome at 1.5-1.7 Mbp ^41,42^.

**Figure 1.**
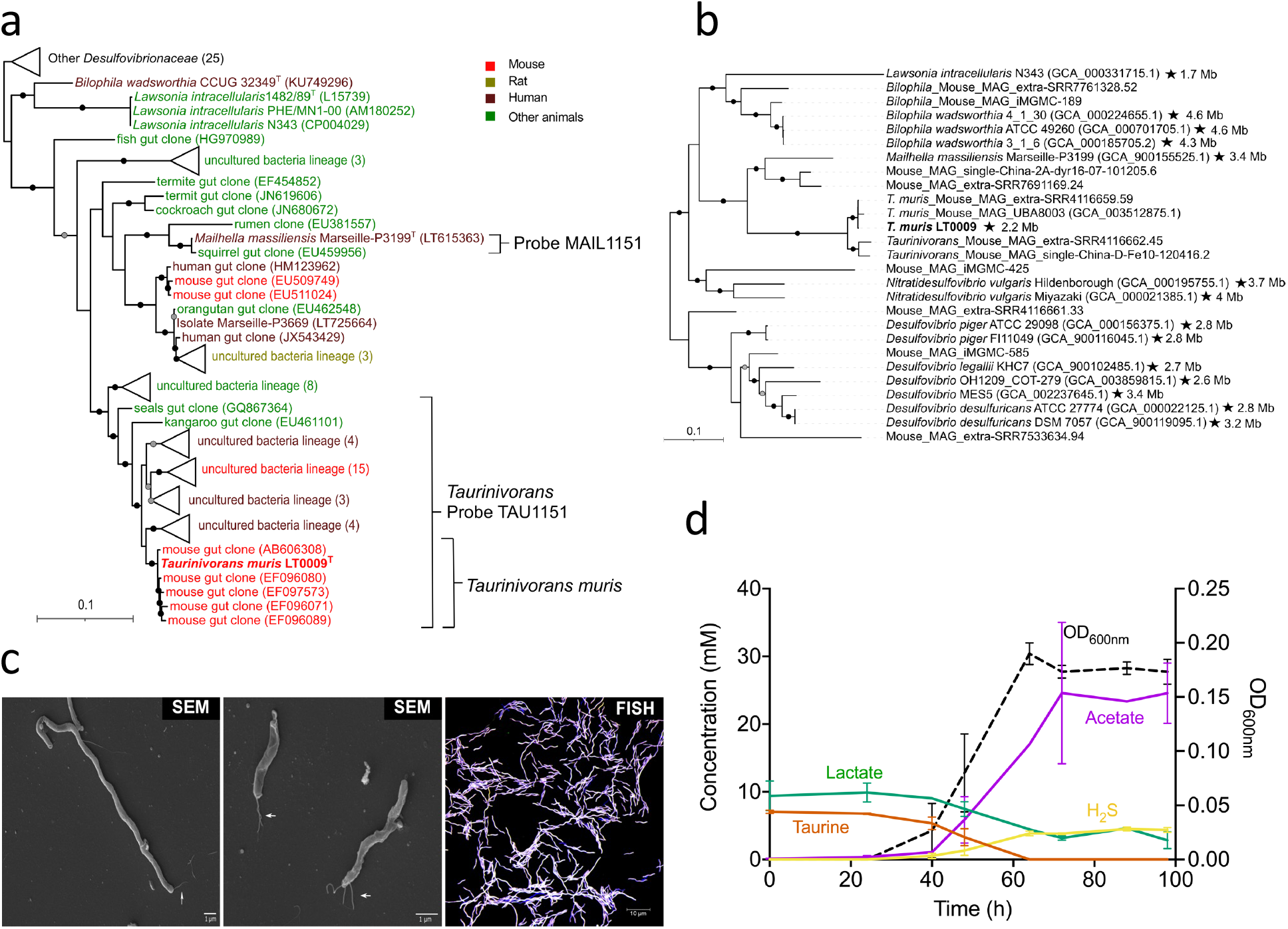
Phylogeny and morphology of the mouse gut-derived taurine-respiring strain LT0009 that represents the new genus/species *Taurinivorans muris* in the family *Desulfovibrionaceae*. **a**. 16S rRNA gene tree and FISH probe coverage. Maximum likelihood branch supports (1000 resamplings) equal to or greater than 95% and 80% are indicated by black and grey circles, respectively. The scale bar indicates 0.1 estimated substitutions per residue. Accession numbers are shown in parentheses. Strain LT0009 is shown in bold and the type strains are marked with a superscript ‘T’. The sequence sources are indicated with different colors (Supplementary Table 2). Sequences were assigned to *Taurinivorans* and *Taurinivorans muris* based on the genus-level similarity cutoff of 94.5% and species-level similarity cutoff of 98.7% ^96^, respectively. The perfect-match coverage of probes TAU1151 for *Taurinivorans* and MAIL1151 for *Mailhella* is indicated. **b**. Phylogenomic tree. Ultrafast bootstrap support values equal to or greater than 95% and 80% for the maximum likelihood tree are indicated with black and grey circles, respectively. Accession numbers are shown in parentheses. Strain LT0009 is shown in bold. Strains with complete genomes (genome size is indicated) are marked with a star. Genomes were assigned to *Taurinivorans* based on the genus-level AAI cut-off value of 63.4% ^97^. The scale bar indicates 0.1 estimated substitutions per residue. **c**. Morphology of LT0009 cells in pure culture. SEM: Scanning electron microscopy images of cells of varying lengths. White arrows indicate the flagella. FISH: Cells hybridized with Cy3-labeled probe TAU1151 and Fluos-labeled probe EUB338mix and counterstained by DAPI. **d**. Growth of strain LT0009 in modified *Desulfovibrio* medium confirmed complete utilization of taurine as electron acceptor concomitant with nearly stoichiometric production of sulfide. Electron donors L-lactate and pyruvate were provided in excess, and their utilization also contributed to acetate formation. Pyruvate in the medium and ammonia released from deamination of taurine were not quantified in this experiment. Lines represent averages of measures in triplicate cultures. Error bars represent one standard deviation.

The Gram-staining of *T. muris* LT0009 was negative. FISH imaging of the LT0009 pure culture with the newly-developed genus-specific 16S rRNA-targeted probe TAU1151 showed cells with a conspicuous spiral-shaped morphology and considerably varying lengths (1.7 to 28 µm) (Fig. 1c). SEM imaging further indicated that LT0009 cells have multiple polar flagella and are thus motile (Fig. 1c).

Complete utilization of taurine as electron acceptor in modified *Desulfovibrio* liquid medium with electron donors lactate and pyruvate in excess, resulted in production of nearly quantitative amounts of H_2_S (Fig. 1d). Strain LT0009 in pure culture did not grow in absence of 1,4-naphthoquinone and yeast extract, indicating an absolute requirement of menaquinone (vitamin K2) and other essential growth factors, respectively. Both growth rate and final growth yields were increased when taurine was provided at 20 and 40 mmol/l concentration in comparison to 10 mmol/l, while growth was inhibited at concentrations ≥60 mmol/l taurine (Supplementary Fig. 3a). Strain LT0009 grew with a lower growth rate and final growth yield when pyruvate was omitted as additional electron donor (Supplementary Fig. 3b). Strain LT0009 grew equally well at a pH range of 6 to 8.5 (Supplementary Fig. 3c) and temperatures between 27-32°C, but with reduced final growth yield at 37 and 42°C (Supplementary Fig. 3d). No colony formation was observed on agar plates under aerobic conditions, suggesting a strict anaerobic lifestyle of *T. muris* LT0009.

### Sulfur and energy metabolism of *T. muris* LT0009

Based on a genome-inferred metabolism prediction of strain LT0009 (Fig. 2a, Supplementary Table 6), we tested its growth with substrates that could serve as energy and sulfur sources in the gut. Fermentative growth with only pyruvate or only taurine, i.e., as both electron donor and acceptor, was not observed. In addition to pyruvate and lactate, LT0009 also used formate as electron donor for growth under taurine-respiring conditions, albeit with an extended lag phase and a lower growth yield. For the alternative electron acceptors tested, LT0009 used 3-sulfolactate and thiosulfate in combination with lactate and pyruvate, but did not grow with 2,3-dihydroxypropane-1-sulfonate (DHPS), isethionate, cysteate, and not with inorganic sulfate or sulfite (Fig. 2b).

**Figure 2.**
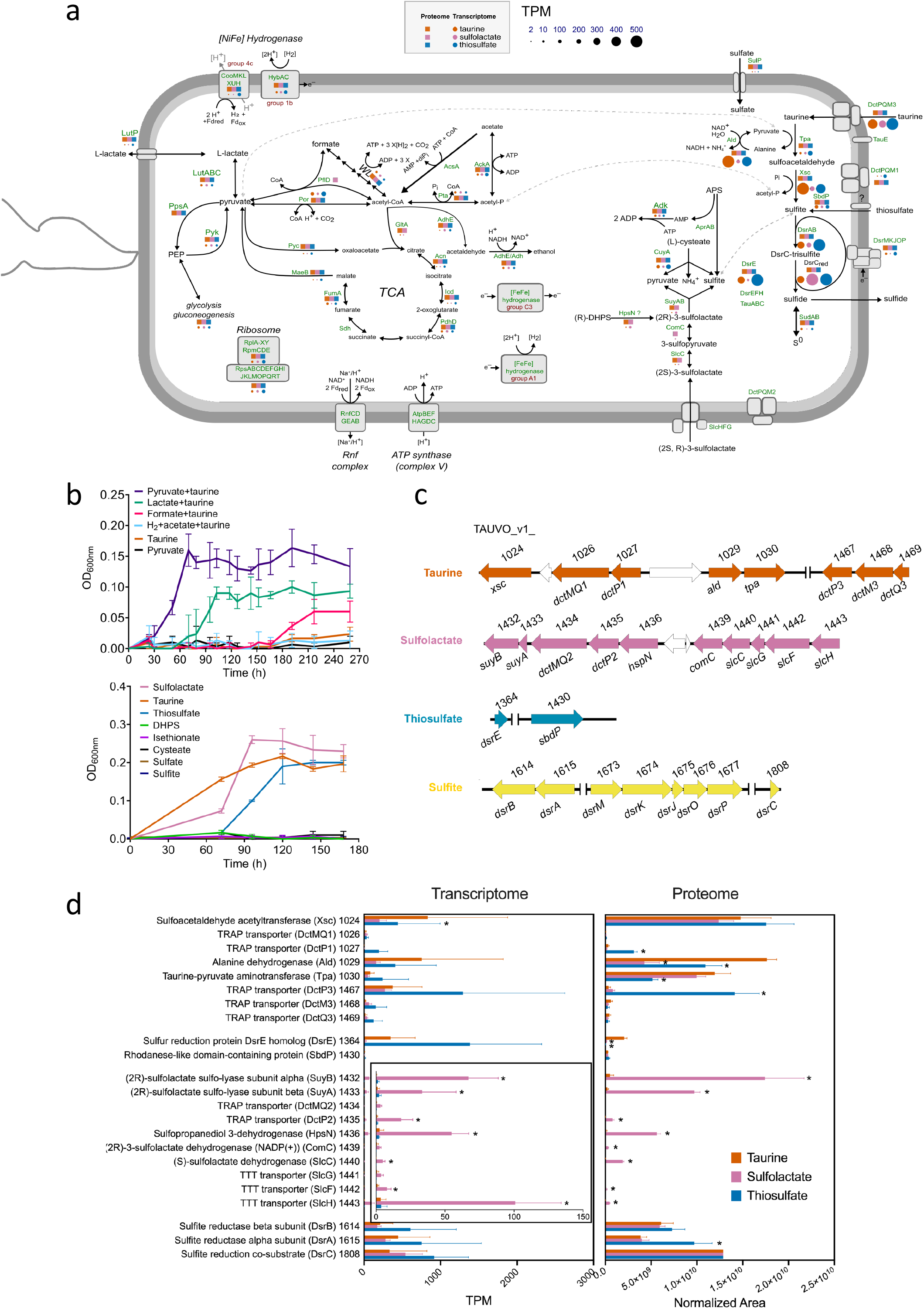
Sulfur-based energy metabolism of *Taurinivorans muris* LT0009. **a**. Cell cartoon of the central sulfur and energy metabolism of LT0009 as determined by genome, transcriptome, and proteome analyses. Genes/proteins detected in the transcriptome and proteome of LT0009 grown with taurine, sulfolactate or thiosulfate as electron acceptor are shown by colored circles and squares, respectively. Circle size indicates gene transcription level normalized as TPM. Proteins of all transcribed genes that are shown were also detected in the proteome, with the exception of AprAB, TauA (TAU_v1_0027, TAU_v1_1344), TauB, TauC, TauE, DctMQ2, SlcG, DsrEFH, AscA, two [FeFe] hydrogenases (TAU_v1_1126, TAU_v1_1901), cytochrome *c* and Sdh. Protein complexes (e.g. Rnf, SlcFGH, DctPMQ2, Atp) are not shown with transcriptome and proteome data because one or more of the complex units were not expressed. The gene annotations are listed in Supplementary Table 6. **b**. Anaerobic growth tests of strain LT0009 with various substrates. Upper panel: All substrates were added at 10 mmol/l concentration, except acetate (20 mmol/l), which was added as carbon source together with H_2_. Lower panel: The different sulfur compounds were added at 10 mmol/l concentration together with pyruvate, lactate, and 1,4-naphthoquinone. OD_600_: optical density at 600 nm. **c**. Organization of sulfur metabolism genes in the LT0009 genome. Numbers show the RefSeq locus tag with the prefix TAUVO_v1. **d**. Comparative transcriptome and proteome analysis of LT0009 grown with lactate and taurine, sulfolactate or thiosulfate as electron acceptor; each in triplicate culture. Numbers following protein names refer to RefSeq locus tag numbers (prefix TAUVO_v1). Protein expression was normalized to DsrC for each growth condition. Bars represent averages of triplicate measures with error bars representing one standard deviation. Asterisk indicates significant (*p*<0.05) differences in gene transcription/protein expression compared to growth with taurine. TCA, tricarboxylic acid cycle; WL, Wood-Ljungdahl pathway; PEP, phosphoenolpyruvate; DHPS, 2,3-dihydroxypropane-1-sulfonate; TPM, transcripts per million.

The metabolic pathways used for growth by respiration with taurine, sulfolactate or thiosulfate and with lactate/pyruvate as electron donors were further analyzed by differential transcriptomics and proteomics. This demonstrated that taurine is degraded via the Tpa-Xsc pathway and the produced sulfite respired via the DsrAB-DsrC pathway (Fig. 2c,d). Pyruvate-dependent taurine transaminase Tpa catalyzes initial conversion of taurine to alanine and sulfoacetaldehyde ^31^. Oxidative deamination of alanine to pyruvate is catalyzed by alanine dehydrogenase Ald (Fig. 2a,c,d). Lack of *sarD* and *islAB* and an inability to grow with isethionate showed that LT0009 does not have the taurine degradation pathway of *B. wadsworthia* ^*26*^. Instead, sulfoacetaldehyde is directly desulfonated to acetyl-phosphate and sulfite by thiamine-diphosphate-dependent sulfoacetaldehyde acetyltransferase Xsc ^29–31^. The acetyl-phosphate is then converted to acetate and ATP by acetate kinase AckA. Strain LT0009 seems to lack candidate genes for the TauABC taurine transporter ^43^. While homologs of *tauABC* are encoded in the genome, the individual genes do not form a gene cluster like in *Escherichia coli* ^*44*^ and were not expressed during growth on taurine (Supplementary Table 7). Instead, the LT0009 genome encodes three copies of gene sets for tripartite ATP-independent periplasmic (TRAP) transporter ^45^ that are co-encoded in the taurine degradation gene cluster and were expressed during growth with taurine, thus are most likely involved in taurine import, including DctPQM1 (with fused DctQM1) (TAUVO_v1_1026 and 1027) and DctPQM3 (TAUVO_v1_1467-1469) (Fig. 2c,d).

(2S)-3-sulfolactate is degraded by LT0009 via the SlcC-ComC-SuyAB pathway as shown by differential expression of these enzymes in cells grown with racemic sulfolactate (Fig. 2c, d, Supplementary Table 7). The dehydrogenases (S)-sulfolactate dehydrogenase SlcC and (R)-sulfolactate oxidoreductase ComC isomerize (2S)-3-sulfolactate to (2R)-3-sulfolactate via 3-sulfopyruvate. (2R)-3-sulfolactate is desulfonated to pyruvate and sulfite by sulfo-lyase SuyAB. The neighboring gene clusters *slcHFG-slcC-comC* and *hpsN-dctPQM-suyAB* were both significantly upregulated in the transcriptome of sulfolactate-grown cells (Fig. 2d). The DctPQM2 (with fused DctQM2) (TAUVO_v1_1434 and 1435) TRAP transporter and the SlcGFH tripartite tricarboxylate transporters (TTT) ^45,46^ are putative sulfolactate importers. The gene cluster further includes a homolog to *hpsN*, encoding sulfopropanediol-3-dehydrogenase ^47^. This enzyme converts (R)-DHPS to (R)-sulfolactate during aerobic catabolism of DHPS by diverse bacteria in soils ^47^ and the ocean ^48^. However, LT0009 did not grow with racemic DHPS when tested (Fig. 2b). The *hpsN* gene was transcribed in LT0009 with taurine, sulfolactate, and thiosulfate treatments, but the HpsN protein was not detected (Supplementary Table 7). LT0009 did not grow with cysteate as electron acceptor under the conditions we used, although it encodes a homolog of L-cysteate sulfo-lyase CuyA that desulfonates L-cysteate to pyruvate, ammonium, and sulfite ^49,50^ (Fig. 2b). The *cuyA* gene was transcribed in the presence of taurine, sulfolactate, and thiosulfate. Furthermore, CuyA was significantly higher expressed in LT0009 with taurine (P<0.001) compared with sulfolactate and thiosulfate (Supplementary Table 7), yet its physiological role in LT0009 remains unclear.

Strain LT0009 respired thiosulfate, such as *B. wadsworthia* strain RZATAU ^51^, but lacks genes for PhsABC thiosulfate reductase ^52^ and thiosulfate reductase from *Desulfovibrio* (EC 1.8.2.5) ^53^, which both (*i*) disproportionate thiosulfate to sulfide and sulfite and (*ii*) are present in the human *B. wadsworthia* strains ATCC 49260, 4_1_30, and 3_1_6. LT0009 has a gene for a homolog of rhodanese-like, sulfur-trafficking protein SbdP (TAUVO_v1_1430) (Supplementary Fig. 4d) ^54^. Rhodaneses (EC 2.8.1.1.) can function as thiosulfate sulfurtransferase and produce sulfite ^55^. Homologs of SbdP are broadly distributed in members of the *Desulfovibrio-Bilophila-Mailhella-Taurinivorans* clade and other *Desulfovibrionaceae* (Supplementary Fig. 4d). The SbdP-homolog in LT0009 (TAUVO_v1_1430) could (*i*) provide sulfite for reduction and energy conservation by the Dsr sulfite reductase system and (*ii*) transfer the second sulfur atom from thiosulfate to an unknown acceptor protein/enzyme. A candidate sulfur-accepting protein is encoded by a *dsrE*-like gene in LT0009 (TAUVO_v1_1364) (Supplementary Fig. 4b). High expression of rhodanese-like sulfur transferases, a DsrE-like protein, and DsrAB sulfite reductase was reported for thiosulfate-respiring *Desulfurella amilsii* ^*56*^. However, the functions of the SbdP-sulfur transferase and the DsrE-like protein in the thiosulfate metabolism of LT0009 remain unconfirmed. First, these proteins are not homologous to the highly expressed *D. amilsii* proteins. Second, comparative transcriptome and proteome analyses were inconclusive as only the transcription of the *dsrE*-like gene was upregulated in LT0009 grown with thiosulfate (Fig. 2d, Supplementary Table 7).

Additional genes homologous to known sulfur metabolism genes whose functions in LT0009 remain enigmatic include *sudAB*, which encode sulfide dehydrogenase for reduction of sulfur or polysulfide to H_2_S ^57^, and *dsrEFH*, which are involved in sulfur atom transfer in sulfur oxidizers (Supplementary Fig. 4c) ^58^.

LT0009 encodes an incomplete pathway for dissimilatory sulfate reduction. While homologs of genes for the sulfate transporter SulP ^59^ and adenylyl-sulfate reductase AprAB were present, absence of genes for sulfate adenylyltransferase Sat and the electron-transferring QmoAB complex (Supplementary Table 6) was consistent with the inability of LT0009 to respire sulfate. Although externally supplied sulfite did not support growth, the DsrAB-DsrC dissimilatory sulfite reductase system was highly expressed in cells grown with taurine, sulfolactate, or thiosulfate (Fig. 2a, d, Supplementary Table 7). This suggests that intracellularly produced sulfite is respired to sulfide via the DsrAB-DsrC system, which includes transfer of electrons from the oxidation of electron donors via the membrane quinone pool and the DsrMKJOP complex (Fig. 2a) ^60^.

Genome reconstruction of LT0009 suggested the potential to utilize lactate, pyruvate, and H_2_ as electron donors (Supplementary Information). We experimentally confirmed that lactate and pyruvate but also formate are used as electron donors for taurine respiration (Fig. 2b).

Notably, *T. muris* employs similar electron acceptors and donors as *B. wadsworthia*, yet they differ in the metabolic pathways to use them.

### Distinct distribution patterns of *Taurinivorans muris* and *Bilophila wadsworthia* suggest different host preferences

*T. muris* is the first taurine-utilizing, sulfidogenic isolate from the mouse gut. *B. wadsworthia* was repeatedly reported as a taurine-degrading member of the murine intestine based on molecular surveys ^23,25,61^. We performed a meta-analysis to compare the presence and relative abundance of *B. wadsworthia-*related and *T. muris*-related sequences across thousands of 16S rRNA gene amplicon datasets from the intestinal tract of diverse hosts. *T. muris*-related 16S rRNA gene sequences were most often detected in the mouse gut, i.e. in 14.4% of all mouse amplicon datasets, but also present in the datasets from multiple other hosts (shrimp, pig, rat, chicken, fish, cow, humans, termites, and other insects) (Fig. 3a, Supplementary Table 8). In comparison, *B. wadsworthia-*related sequences are most widespread in the human gut, i.e. in 30.7% of human gut amplicon datasets, but are also prevalent in pig (15.8%), chicken (13.7%), and rat (9.8%) and occasionally detected in other hosts (Fig. 3a). We also identified *B. wadsworthia-*related sequences in 7.5% of mouse datasets. Notably, *T. muris*- and *B. wadsworthia-*related sequences co-occurred only in 28 mouse datasets, which suggests competitive exclusion possibly due to competition for taurine. Furthermore, we found that 82% of the *B. wadsworthia*-positive samples are from mice that were ‘humanized’ by receiving human feces transplants or human strain consortia ^62–65^, which suggests a much lower prevalence of *B. wadsworthia* strains that are indigenous in mice. *T. muris*-related sequences represented >5% of the total community in 2.8% of mouse gut datasets (Fig. 3a). Such very high relative 16S rRNA gene abundances were more often observed in mice on high-fat diets ^66,67^, but sporadically also in mice on standard chow and other diets (Fig. 3b) ^68^.

**Figure 3.**
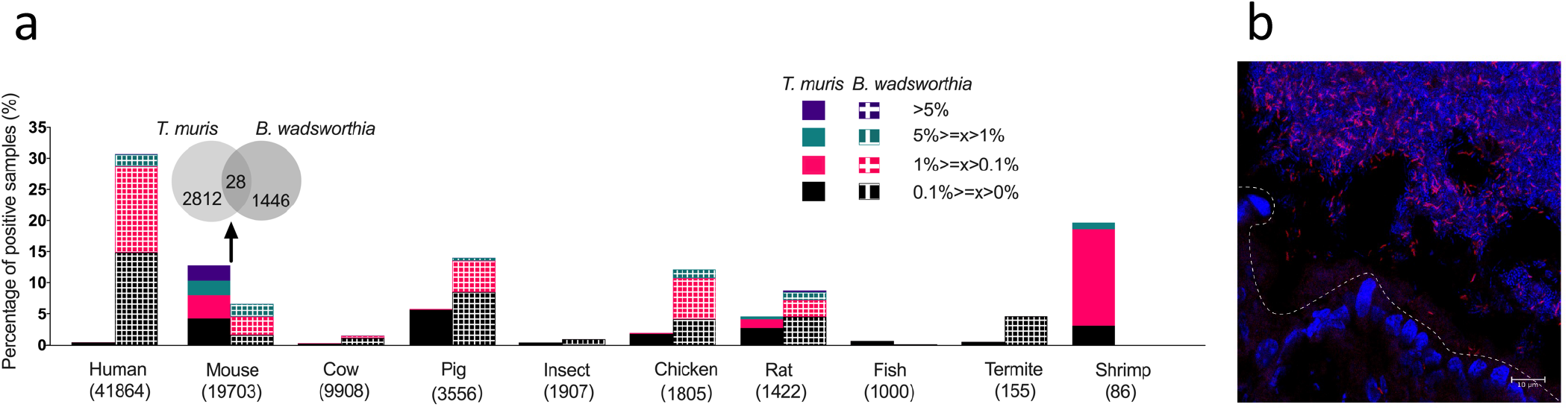
*Taurinivorans muris* and *Bilophila wadsworthia* have distinct host distribution patterns. **a**. Occurence and prevalence of *Taurinivorans muris*- and *Bilophila wadsworthia*-related sequences in 16S rRNA gene amplicon datasets of human and animal guts. *T. muris*- and *B. wadsworthia*-like sequences at 97% similarity cut-off are expressed as percentages of positive samples in each host (the numbers of samples used for the analysis are shown in parenthesis) and different colors indicate percentages of samples positive for *T. muris* and *B. wadsworthia* at different relative abundance ranges. Hosts with less than 20 amplicon samples are not shown. *T. muris-* and *B. wadsworthia-*related sequences co-occur in only 28 mouse gut samples as shown by the Venn diagram. **b**. Visualization of *Taurinivorans* in a colon tissue section of a mouse fed a polysaccharide- and fiber-deficient diet ^98^ by FISH. TAU1151-Cy3-labeled *Taurinivorans* cells appear in pink and the remaining bacterial cells and tissue in blue due to DAPI-staining. The dashed line indicates the border between epithelial cells and gut lumen.

Overall, *T. muris* is considerably more abundant and prevalent in the mouse gut microbiome than *B. wadsworthia*. Notably, a mouse native *B. wadsworthia* strain has not yet been isolated. Our phylogenomic analysis of all de-replicated, high-quality *Desulfovibrionaceae* MAGs from the integrated mouse gut metagenome catalog (iMGMC) ^69^ revealed two MAGs form a well-supported monophyletic group with *B. wadsworthia* strains (Fig. 1b). Mouse MAG iMGMC-189 has a minimum ANI of 82% and AAI of 79% to *B. wadsworthia*, which suggests it represents the population genome of a new, murine *Bilophila* species. Mouse MAG extra-SRR7761328.52 is more distantly related and has a minimum ANI of 78% and AAI of 65% to *B. wadsworthia*. Both mouse MAGs encode the taurine degradation pathway (*tpa-sarD-islAB*) of *B. wadsworthia* (Supplementary Fig. 2), while the pathway for sulfolactate degradation (*slcC-comC-suyAB*) is absent in MAG extra-SRR7761328.52.

In general, genes for utilization of diverse organosulfonates are widely and patchily distributed in the *Desulfovibrio-Bilophila-Mailhella-Taurinivorans* clade (Supplementary Fig. 2) ^33^. Other mouse *Desulfovibrionaceae* that encode the capability for taurine respiration include (i) the *Desulfovibrio*-affiliated MAGs extra-SRR7533634.94 and iMGMC-585 with the *tpa-xsc* pathway and (ii) the *Mailhella*-related MAG extra-SRR7691169.24 with the *tpa-sarD-islAB* pathway (Supplementary Fig. 2).

### Taurine degradation is the main *in vivo* realized nutritional niche of *Taurinivorans muris*

We next performed metatranscriptome analysis and re-analyzed published metatranscriptome datasets of gut samples from different mouse models to reveal the metabolic pathways that are most expressed by *T. muris* in its murine host.

In our gnotobiotic model, strain LT0009 or a mock control were added to germ-free mice stably colonized with the synthetic OMM^12^ community (Fig. 4a). Strain-specific qPCR assays showed that ten OMM^12^ strains and strain LT0009 colonized the mice (Fig. 4b). Consistent with previous studies, strains *A. muris* KB18 and *B. longum* subsp. animalis YL2 were not detected ^35,70^. Colonization of LT0009 did not affect the abundance of other strains, which indicated that LT0009 occupied a free niche in the intestinal tract of this gnotobiotic mouse model. The taurine metabolism (*tpa, ald, xsc*) and sulfite reduction (*dsrAB, dsrC*) genes were in the top 5% expression rank of all LT0009 genes (Fig. 4c). In contrast, gene expression of the putative thiosulfate transferase (*sbdP*) ranked at 17% and of sulfolactate degradation (*suyAB, slcC, comC*) ranked from 62% to 88% of all LT0009 genes (Fig. 4c). *T. muris* LT0009 thus largely occupied the vacant taurine-nutrient niche in the intestinal tract of OMM^12^ mice.

**Figure 4.**
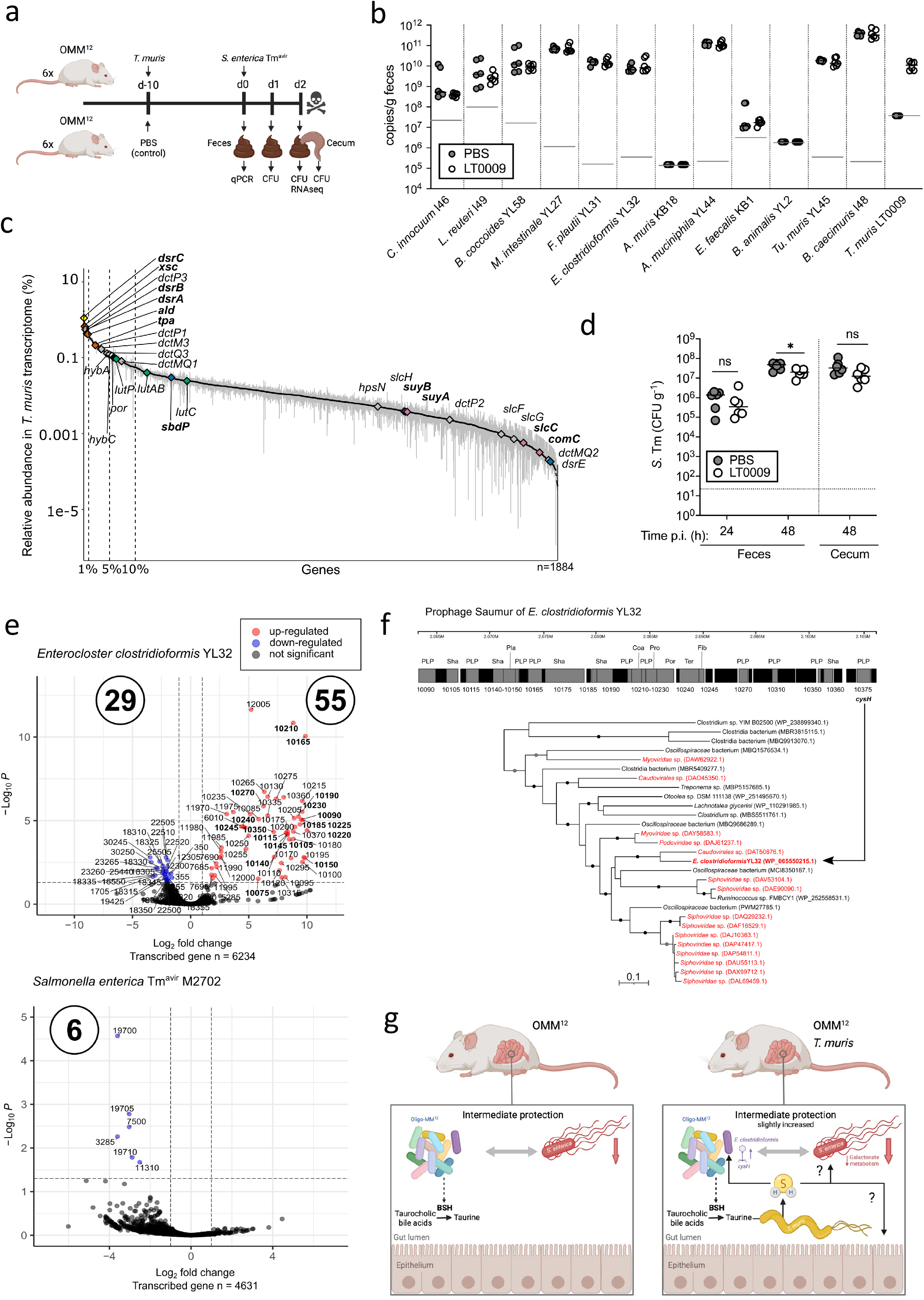
*Taurinivorans muris* mainly respires taurine *in vivo* and slightly enhances colonization resistance against *Salmonella enterica* in a gnotobiotic mouse model. **a**. Schematic outline of the gnotobiotic mouse experiment. Mice stably colonized with the 12-strain Oligo-Mouse-Microbiota (OMM^12^) were inoculated with *T. muris* LT0009 (n=6) or sterile phosphate-buffered saline (PBS) as control (n=6) and, after 10 days, orally and rectally infected with *S. enterica* Tm^avir^ M2702. Mice were sacrificed two days post infection (p.i.). Fecal samples were used for strain-specific 16S rRNA gene-targeted quantitative PCR (qPCR). Fecal and cecal samples at 24 h and 48 h p.i. were used for analysis of colony forming units (CFU) of *S. enterica* Tm. Fecal samples at 48 h p.i. were used for metatranscriptomics (RNAseq). **b**. Absolute abundances (16S rRNA gene copy numbers per gram feces) of each OMM^12^ strain and strain LT0009 on day 10 in feces of mice with LT0009 and the PBS-control mice. Small horizontal lines indicate median values. Gray horizontal lines indicate the detection limit of each strain-specific qPCR assay. **c**. Ranked relative transcript abundance of LT0009 genes in OMM^12^ mice fecal metatranscriptomes. Each point is the mean relative abundance of a gene and error bars correspond to the 95% confidence interval of the mean (n = 3). The total number of transcribed LT0009 genes is shown (n=1884). Genes for taurine (*tpa, xs*c, *ald*), sulfite (*dsrAB, dsrC*), sulfolactate (*suyAB, sclC, comC*) thiosulfate (*sbdP, dsrE*), pyruvate (*por*), lactate (*lutABC*), and hydrogen (*hybA, hybC*) metabolism are shown in different colors. Sulfur metabolism genes are further highlighted in bold font. Vertical dashed lines delineate the top 1%, 5%, and 10% expression rank of all protein-coding genes in the LT0009 genome (n=2059). **d**. CFU of *S. enterica* Tm at 24 h and 48 h p.i. in the feces and at 48 h p.i. in the cecal content. Small horizontal lines indicate median values. The dotted horizontal line shows the CFU detection limit. The asterisk indicates significant differences (p<0.05; ANOVA using Kruskal-Wallis and Dunn’s Multiple Comparison test) between *S. enterica* Tm^avir^ CFU in mice with LT0009 and the PBS-control mice. ns, not significant. **e**. Volcano plots of differential gene transcription of *S. enterica* Tm^avir^ M2702 and *E. clostridioformis* YL32 in OMM^12^ mice with and without LT0009. The x-axis shows log-fold-change in transcription and the y-axis shows the negative logarithm10-transformed adjusted p values. Blue dots show significantly down-regulated genes (adjusted p-value <0.05, log2 fold change <-1) in mice with LT0009 and are labeled with locus tag numbers. Up-regulated *E. clostridioformis* YL32 prophage genes in I5Q83_10075-10390 are highlighted in bold. **f**. Structure of the activated prophage gene cluster of *E. clostridioformis* YL32 and phylogeny of its encoded phosphoadenosine-phosphosulfate reductase (CysH). Virus- and bacteria-encoded sequences are shown in red and black, respectively. The maximum likelihood CysH tree is midpoint rooted. Ultrafast bootstrap support values equal to or greater than 95% and 80% for the maximum likelihood tree are indicated with black and grey circles, respectively. The identity of the 61 genes in the prophage region (I5Q83_10075-10390) of *E. clostridioformis* YL32 as predicted by PHASTER ^91^. Genes encoding hypothetical proteins are in black and annotated genes are in grey. Numbers indicate the locus tag. PLP, phage-like protein; Sha, tail shaft; Pla, plate protein; Coa, coat protein; Pro, protease; Por, portal protein; Ter, terminase; Fib, fiber protein. **g**. Cartoon illustrating the impact of *T. muris* LT0009 on *S. enterica* Tm^avir^ M2702 colonization resistance in the OMM^12^ mouse model. Created with Biorender.com.

Metatranscriptome analysis of intestinal samples from conventional laboratory mice on various diets (e.g. high-glucose; high-fat/low-carbohydrate; low-fat/high-carbohydrate) and with different genetic backgrounds (wildtype; plin2) also showed that taurine degradation and sulfite respiration were within the top 5% of expressed LT0009 genes (Supplementary Fig.5). Collectively, this demonstrated that taurine is the predominant electron acceptor for energy conservation of *T. muris* in the murine intestinal tract.

Free taurine in the murine intestine largely derives from microbial deconjugation of host-derived taurocholic bile acids ^71,72^. LT0009 does not encode genes for bile salt hydrolase (BSH) and is thus likely dependent on other gut microbiota members for liberation of taurine from taurocholic bile acids. BSH genes are encoded across diverse bacterial taxa in the human and mouse gut ^71,73,74^. In agreement with previous studies of bile acid transformations in the OMM^12^ model ^75,76^, we identified BSH genes in seven OMM^12^ strains (Supplementary Table 5). The expression of BSH genes in these strains did not change significantly with the presence of LT0009. Yet, BSH gene transcription increased in *E. clostridioformis* YL32, *E. faecalis* KB1, *B. caecimuris* I48, and *M. intestinale* YL27, and decreased in *L. reuteri* I49 (Supplementary Table 5). Bile acid deconjugation by some of these OMM^12^ strains has been confirmed *in vitro* ^*76*^. Specifically, *B. caecimuris* I48, *B. animalis* YL2, *E. faecalis* KB1, and *M. intestinale* YL27 were tested positive for deconjugation of taurine-conjugated deoxycholic acid, while *E. clostridioformis* YL32 was either tested negative or inhibited by addition of the bile acids. The down-regulation of the BSH gene in *L. reuteri* I49 is consistent with the *in vitro* deconjugation capacity of this strain for glycine-conjugated deoxycholic acid but not for taurine-conjugated deoxycholic acid ^76^ and the generally lower abundance of glycine-conjugated bile acids in rodents ^72^. We hypothesize that taurine degradation by LT0009 could provide a selective feedback mechanism on expression of BSHs for taurocholic bile acid deconjugation in the OMM^12^ model.

Thiosulfate is presumably a constantly present electron acceptor for microbial respiration in the mammalian gut as it is generated by mitochondrial H_2_S oxidation in the gut epithelium ^1^. The sulfide oxidation pathway is mainly located apically in the crypts of human colonic tissue at the interface to the gut microbiota ^77^. The amount of thiosulfate supplied into the gut lumen will depend on epithelial H_2_S metabolism ^1,77^. The expression level of the putative SbdP thiosulfate transferase of *T. muris* ranked relatively high with 15-24% of all LT0009 genes across all mouse gut samples (Fig. 4c, Supplementary Fig. 5). However, the function of this protein remains unconfirmed as its expression was not differentially upregulated in the thiosulfate-metabolizing *T. muris* pure culture (Fig. 2a,d, Supplementary Table 7).

*In vivo* taurine respiration, and potentially thiosulfate respiration, are likely fueled by pyruvate, H_2_, and lactate as electron donors, as expression of genes for their oxidation ranked at 5-7% (*por*), 1.5-9.1% (*hybAC*), and 2.5-31% (*lutABC, lutP*) of all LT0009 genes across all mouse gut metatranscriptomes, respectively (Fig. 4c, Supplementary Fig. 5).

### *Taurinivorans muris* LT0009 slightly increased colonization resistance against *S. enterica* and activated a sulfur metabolism gene-encoding prophage in a gnotobiotic mouse model

The human enteropathogen *S. enterica* Tm can invade and colonize the intestinal tract by utilizing various substrates for respiratory growth that are available at different infection stages ^78^. The gnotobiotic OMM^12^ mouse model provides intermediate colonization resistance against *S. enterica* Tm ^35^ and is widely used as a model system of modifiable strain composition for investigating causal involvement of cultivated mouse microbiota members in diverse host diseases and phenotypes ^79^. Yet, a bacterial isolate from the mouse gut with proven dissimilatory sulfidogenic capacity was not available until now ^80^. *T. muris* has fundamental physiological features that could on the one hand contribute to colonization resistance against *S. enterica* Tm by direct competition for pyruvate ^81^, lactate ^82^, H_2_ ^83^, formate ^84^, and host-derived thiosulfate ^84,85^. On the other hand, *T. muris* could also promote *S. enterica* Tm expansion during inflammation by fueling tetrathionate production through enhanced intestinal sulfur metabolism ^18,86^. Furthermore, expansion of sulfidogenic *Deltaproteobacteria* commensals and the *tpa-xsc-dsr* pathway in the metagenome fueled by host-derived taurine was shown to increase protection against the enteropathogen *Klebsiella pneumoniae* in mice; with sulfide-mediated inhibition of aerobic respiration by pathogens being proposed as a generic protective mechanism ^34^. Notably, our re-analysis of the 16S rRNA gene amplicon data from the wildR mouse model of this study identified *T. muris* as the dominant deltaproteobacterium (*Desulfobacterota*) of the protective community (Supplementary Fig. 6). Given that taurine respiration via the sulfidogenic *tpa-xsc-dsr* pathway is the main energy niche of *T. muris* in the mouse gut (Fig. 4c, Supplementary Fig. 5), enhanced resistance in the wildR mouse model was thus likely primarily due to the activity of *T. muris*.

Here, we investigated the impact of *T. muris* LT0009 during the initial niche invasion of *S. enterica* Tm using an avirulent, non-colitogenic strain ^87^. Compared to OMM^12^ mice without LT0009, mice colonized with the OMM^12^ and LT0009 had a slightly reduced load of *S. enterica* Tm at 48 p.i. that was significant in feces but not in cecum (Fig. 4d). Comparative metatranscriptome analysis did not provide evidence for a mechanism of direct interaction as only six *S. enterica* Tm genes were differentially expressed, i.e. significantly downregulated, in the presence of strain LT0009 (Fig. 4e, Supplementary Table 9). Three of the six genes are involved in transport and metabolism of galactonate (D-galactonate transporter DgoT, D-galactonate dehydratase DgoD, 2-dehydro-3-deoxy-6-phosphogalactonate aldolase DgoA), which is produced by some bacteria as an intermediate in D-galactose metabolism and is also present in mammalian tissue and body secretions ^88^. D-galactonate catabolism capability was suggested as a distinguishing genetic feature of intestinal *Salmonella* strains compared to extraintestinal serovars, with serovars Typhi, Paratyphi A, Agona, and Infantis lacking genes for utilizing D-galactonate as a sole carbon source ^89^. The putative D-galactonate transporter DgoT in *Salmonella enterica* serovar Choleraesuis was identified as a virulence determinant in pigs ^90^. The OMM^12^ strains and LT0009 do not encode the DgoTDAKR galactonate pathway. The significance of galactonate for *S. enterica* Tm gut colonization and competition remains to be elucidated.

Colonization of *T. muris* LT0009 in the gnotobiotic mice had variable impact on the differential gene expression pattern of the OMM^12^ members (Fig. 4e, Supplementary Fig. 7 and Table 9). While gene expression was not significantly affected in *L. reuteri* I49, *E. clostridioformis* YL32 was most affected with 84 differentially expressed genes (55 up-regulated and 29 down-regulated) (Fig. 4e). Most of the significantly up-regulated genes (n=41) in *E. clostridioformis* YL32 are clustered in a large genomic region (I5Q83_10075-10390) that encoded various phage gene homologs and was identified as a prophage using PHASTER (Fig. 4f). This prophage, named YL32-pp-2.059, Saumur, is among a set of thirteen previously identified prophages of the OMM^12^ consortium that represent novel viruses, were induced under various *in vitro* and/or *in vivo* conditions, and constitute the temporally stable viral community of OMM^12^ mice ^91^. We show that colonization of LT0009 in the OMM^12^ mouse model selectively enhanced the transcriptional activity of the *E. clostridioformis* YL32 prophage Saumur, which carries a gene (I5Q83_10375) for phosphoadenosine-phosphosulfate reductase (CysH) that functions in the assimilatory sulfate reduction pathway of many bacteria (Fig. 4f). Various organosulfur auxiliary metabolic genes, particularly *cysH*, are widespread in environmental and human-associated viromes, which suggests viruses augment sulfur metabolic processes in these environments, including the gut ^92^. Addition of physiologically relevant concentrations of sulfide to a *Lactococcus lactis* strain culture resulted in increased production of viable particles of its phage P087 ^92^. We thus hypothesize that *T. muris* LT0009 not only impacts intestinal sulfur homeostasis via its sulfur metabolism but also by H_2_S-mediated activation of the sulfur metabolism gene-expressing phage Saumur in *E. clostridioformis* YL32. If activation of prophage Saumur contributes to protection from *S. enterica* Tm remains subject of further study.

Our results suggest that the mouse commensal *T. muris* can enhance colonization resistance against different enteropathogens. In addition to direct inhibition of aerobic respiration of pathogens by H_2_S as shown previously ^34^, indirect resistance mechanisms that are dependent on the respective pathogen and composition of the resident microbiota can be at play. For example, H_2_S could activate prophages and expression of their auxiliary metabolic genes ^92^ and thereby modulate microbiome functions (Fig. 4g).

## Conclusions

Dissimilatory sulfur metabolism with production of H_2_S is a core metabolic capability of the mammalian gut microbiota that is carried out by specialized bacteria ^33,93^. As in humans and other animals, bacteria of the phylum *Desulfobacterota* (formerly *Deltaproteobacteria*), specifically of the *Desulfovibrio-Mailhella-Bilophila* lineage ^37^, are important sulfidogens in the mouse gut and appear to be more abundant in wild mice compared to untreated laboratory mice ^94,95^. However, the diversity of sulfidogenic microorganisms and their metabolic pathways in the intestinal tract of non-human animals, including mice that represent important experimental models, remain insufficiently understood. The first mouse gut-derived *Desulfobacterota* strain (*Desulfovibrio* strain MGBC000161) was recently isolated ^94^. Here, we contribute the sulfidogenic strain LT0009, representing the new genus *Taurinivorans*, to the growing collection of publicly available bacterial strains from the mouse ^80,94^. We further describe the fundamental metabolic properties and realized lifestyle of *Taurinivorans*, which relies on taurine as primary electron acceptor for energy conservation *in vivo* and can contribute to the protective effect of the commensal mouse gut microbiota against enteropathogens (Fig. 5). *T. muris* strain LT0009 is the first murine *Desulfobacterota* isolate with a physiologically proven dissimilatory sulfur metabolism and thus significantly extends the experimental options to study the role of sulfidogenic bacteria in gnotobiotic mouse models.

**Figure 5.**
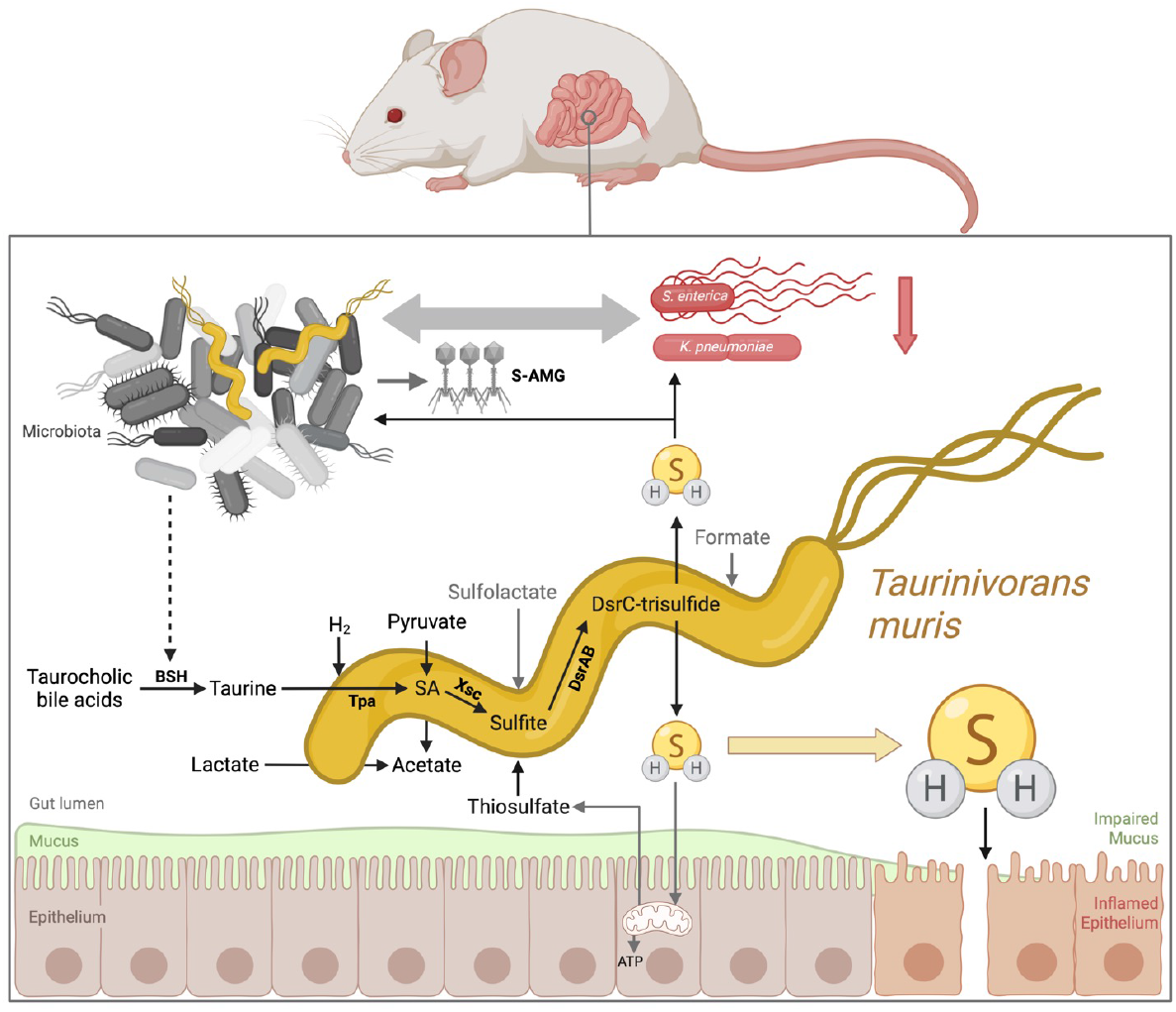
Sulfur energy metabolism and interaction scheme of *Taurinivorans muris* in the mouse gut. *T. muris* mainly utilizes taurine as the main electron acceptor for anaerobic respiration in the gut but is also capable of thiosulfate and sulfolactate respiration. Pyruvate, lactate, and likely hydrogen are the main electron donors of *T. muris*, while formate could also be used. Taurine is cleaved from host-derived taurocholic bile acids by other gut bacteria via bile salt hydrolase (BSH). Thiosulfate derives from mitochondrial oxidation of H_2_S in the gut epithelium. *T. muris* produces H_2_S from taurine via pyruvate-dependent taurine transaminase (Tpa), sulfoacetaldehyde (SA) acetyltransferase (Xsc), and dissimilatory sulfite reductase (DsrAB). H_2_S can have various effects on the gut microbiota and host health. For example, excess H_2_S can impair mucus integrity ^3^. H_2_S can enhance resistance against enteropathogens by directly inhibiting enzymes in aerobically respiring *Klebsiella pneumoniae* ^*34*^. H_2_S could further impact microbial interactions and intestinal metabolism by activating phages and expression of their auxiliary metabolic genes, such as those involved in sulfur metabolism (S-AMG) ^92^. Created with Biorender.com.

## Acknowledgments

We thank Bernhard Schink (University of Konstanz, Germany) for latin naming, Daniela Gruber (Core Facility of Cell Imaging and Ultrastructure Research, University of Vienna) and Isabella Böhm for help with scanning electron microscopy, Markus Schmid for help with FISH imaging, and Jasmin Schwarz and Gudrun Kohl (Joint Microbiome Facility) for sequencing. We also thank Holger Daims, Kerrin Steensen, the DOME gut group members in Vienna as well as our colleagues at University of Konstanz and LMU Munich for fruitful discussions and support. This work was financially supported by the Austrian Science Fund (FWF; project grants I2320-B22 and DOC 69-B), the Deutsche Forschungsgemeinschaft (DFG; grants SCHL1936/3-4, STE 1971/7-1), the Konstanz Research School Chemical Biology (KoRS-CB), and the China Scholarship Council (PhD fellowship grant No. 201606850092).

## Author contributions

AL conceived the study. HY, SB, CE, and BTH performed experiments and analyzed data. BZ generated relevant preliminary data. CWH, BH, and PP provided bioinformatic support. BTH, PP, BS, and DS helped with experimental design and data interpretation. HY and AL wrote the article. All authors discussed the results and revised the manuscript.

## Competing interests

The authors declare no competing interests.

**Correspondence** and requests for materials should be addressed to Alexander Loy.

