## Supplementary Information for "A novel taurine-respiring murine gut bacterium contributes to colonization resistance against enteropathogens"

**Content**

Supplementary text

*Materials and methods*

*Results & discussion*

Supplementary table legends

Supplementary figures and legends

Supplementary references

#### Supplementary text

##### Materials and methods

**Enrichment and isolation of strain LT0009**

Intestinal content (cecum and colon) of wild-type C57BL/6 mice co-housed with access to normal chow and water *ad libitum* was used as inoculum for the enrichment cultures. Gut content was collected immediately after sacrifice of the mice in an anaerobic chamber and homogenized in 50 ml sterile, anoxic 30 mmol/l bicarbonate buffer (pH 7) for inoculation. A modified *Desulfovibrio* medium (based on DSMZ medium 641, www.dsmz.de) in which all sulfur-containing chemicals were omitted (i.e., $Na_2SO_4$, $Na_2S_2O_3$, $MgSO_4$, and $Na_2S$) was used for enrichment of taurine-respiring microorganisms in presence of lactate and pyruvate as electron donors. The basal medium consisted of (per liter of final medium) 1 g $NH_4Cl$, 2.1 g $Na_2Cl$, 0.825 g $MgCl_2$ x $6H_2O$, 0.1 g $CaCl_2$ x $2H_2O$, 0.5 g $KH_2PO_4$, 1 g yeast extract (Thermo Fisher Scientific, US), 1 mL trace element solution SL-10 (DSMZ medium 320), and 1 ml selenite-tungstate solution (DSMZ medium 385). The basal medium was autoclaved, supplemented with 0.2 µm filter-sterilized $NaHCO_3$ stock solution (2 g $NaHCO_3$ per liter of final medium) and 10 ml vitamin solution (DSMZ medium 141), and placed in an anaerobic chamber (Coy labs, USA) under anoxic atmosphere (85% $N_2$, 10% $CO_2$, 5% $H_2$) for two days to ensure sufficient equilibration. All further work was done in the anaerobic chamber if not otherwise stated. The final modified *Desulfovibrio* medium was supplemented by (final concentrations) 10 mmol/l taurine, 10 mmol/l L-lactate, 20 mmol/l pyruvate, and 0.2 mg/l 1,4-naphthoquinone; each added from filter-sterilized stock solutions that were prepared in into autoclaved serum bottles and crimp sealed with sterile rubber stoppers and made anoxic by sparging with $N_2$ gas for 0.5 h. Four ml of this final medium were distributed into sterile 20 ml Hungate tubes closed with sterile butyl rubber stoppers and inoculated each with one ml of supernatant of the homogenized intestinal content. The cultures were incubated at 37°C, periodically sampled for $H_2S$ and taurine quantification, and sub-cultivated into new tubes by transfer of one ml into four ml fresh media every two to three days over 8 weeks. Isolation of enriched taurine-metabolizing, sulfidogenic microorganisms was performed using differential agar plates in the anaerobic chamber. The plates contained the basal medium supplemented with 2.5 g/l Na-L-lactate, 1.25 g/l taurine, 2.2 g/l sodium pyruvate, 200 µg/l 1,4-naphthoquinone, and 0.5 g/l ammonium ferric citrate and were solidified with 1.5% agar; the $H_2S$ produced from taurine and the ferric iron form a black iron sulfide precipitate. Black colonies were picked and streaked onto fresh plates until a uniform colony morphology was observed. Single colonies were inoculated back into 5 ml liquid medium and these cultures were further purified by dilution to extinction, yielding strain LT0009 as an isolate. The isolation process was monitored by microscopy and direct Sanger sequencing of bacterial 16S rRNA gene amplicons. The purity of the culture was additionally confirmed by fluorescence *in situ* hybridization (FISH) using a newly developed LT0009-specific probe. The isolate was stored in 5% DMSO (1 ml culture plus 1 ml 10% DMSO in sterile water at -80°C.

**Growth experiments**

Strain LT0009 was routinely grown at 37°C in 5 ml modified *Desulfovibrio* medium described above in 20 ml Hungate tubes under an atmosphere of 10% $CO_2$, 5% $H_2$, and 85% $N_2$ and closed with butyl rubber stoppers. Substrate utilization tests were also performed in 96-well plates (transparent, F-bottom, Greiner bio-one cellstar) in the anaerobic chamber. During these incubations, optical density at 600 nm ($OD_{600nm}$) was measured every 30 minutes in a plate reader (Multiskan Go, Thermo Scientific) with 5 s shaking prior to measurements.

To determine the optimal taurine concentration for LT0009 growth, 200 µl of the modified *Desulfovibrio* medium was supplemented with different taurine concentrations (10, 20, 40, 60, 80, and 100 mmol/l, each in triplicate), amended with 10 mmol/l lactate, 20 mmol/l pyruvate, and 0.2 mg/l 1,4-naphthoquinone, and inoculated with 40 µl of LT0009 culture. Growth was measured with the plate reader. Growth of LT0009 in 10 mmol/l taurine was additionally tested with the same setting but without pyruvate.

For substrate utilization tests, the modified *Desulfovibrio* medium was supplemented with different electron donors with taurine as the electron acceptor. Tested substrate combinations included: lactate/taurine, formate/taurine, pyruvate/taurine, hydrogen/acetate/taurine. Fermentative growth with pyruvate or taurine in absence of a dedicated electron donor was also tested. All cultures were amended with 10 mmol/l of each substrate and 1,4-naphthoquinone at 0.2 mg/l. Hydrogen was flushed into Hungate tubes to a pressure of two bars.

Utilization of different organic and inorganic sulfur compounds (each 10 mmol/l) as electron acceptors was tested in modified *Desulfovibrio* medium with a final concentration of 10 mmol/l lactate, 20 mmol/l pyruvate, and 0.2 mg/l 1,4-naphthoquinone. The sulfur compounds included: taurine（Sigma-Aldrich, cat.no. T8691), sodium sulfate (Carl Roth, cat.no. 8560.3), sodium sulfite (Sigma-Aldrich, cat.no. 71922), racemic sulfolactate (synthesized as described previously [1]), sodium thiosulfate (Sigma-Aldrich, cat.no. 217263), L-cysteate (Sigma-Aldrich, cat.no. 30170), racemic 2,3-dihydroxypropane-1-sulfonate (DHPS) (synthesized as described previously [2]), and isethionate (Sigma-Aldrich, cat.no. 820708010). 100 µl of LT0009 culture was transferred into triplicate tubes to test the growth with each sulfur compound. For differential proteomics and transcriptomics, cells of LT0009 were harvested from cultures grown with taurine, sulfolactate, or thiosulfate in the late exponential growth phase.

The pH range for optimal growth was assessed at pH 4.0, 5.0, 6.0, 6.5, 7.0, 7.5, 7.9, and 8.5 at 37°C in triplicate Hungate tubes. The pH was adjusted with 0.1 mol/l HCl or 1 mol/l NaOH followed by filter sterilization. The temperature range for optimal growth was tested at 20, 27, 30, 32, 37, 42, 61, and 75°C in triplicate at pH 7.2. OD at 600 nm was measured periodically during growth using a spectrophotometer (Ultrospec 10, Amersham Bioscience). LT0009 was incubated at 37°C on modified *Desulfovibrio* medium agar plates under aerobic conditions to test its susceptibility to oxygen.

**Substrate and metabolite quantification**

For quantification of substrate removal and metabolite formation, strain LT0009 was grown in triplicate in 20 ml Hungate tubes with an initial concentration of 10 mmol/l taurine, 10 mmol/l lactate, 20 mmol/l pyruvate, and 0.2 mg/l 1,4-naphthoquinone. The culture was subsampled at time points 0, 24, 40, 48, 64, 72, 88, and 98 h for OD and substrate/metabolite quantification. Taurine in cell-free samples was quantified using 4-fluoro-7-nitrobenzofurazan as a derivatizing agent as previously described [3] using an Infinite 200 PRO spectrophotometric microplate reader (TECAN Group Ltd., Männedorf, Switzerland) with excitation set at 470 nm and emission set at 530 nm. $H_2S$ in cell-free samples was quantified spectrophotometrically as previously described [4] with absorbance measurements at 670 nm using an Infinite 200 PRO spectrophotometric microplate reader (TECAN Group Ltd., Männedorf, Switzerland). Dilutions of a sodium sulfide solution (Sigma-Aldrich, cat.no. 71988) were used as external standards for $H_2S$ quantification. Short-chain fatty acids (SCFA) were measured using P/ACE-MDQ capillary electrophoresis equipped with an UV-detector (Beckman Instruments, Krefeld, Germany). Samples were diluted 1:20 with a working solution consisting of 0.01 mol/l NaOH, 0.5 mmol/l $CaCl_2$, and 0.1 mmol/l caproate (internal standard). A dilution series of a stock mixture of sulfate, formate, succinate, acetate, lactate, propionate, butyrate, and valerate (1 mmol/l each) was used as external SCFA standards for quantification. CEofixTM Anions 5 kit (Analis, Belgium) was used for SCFA measurement according to the manufacturer's instruction.

**Electron and fluorescence microscopy**

For scanning electron microscopy, 5 ml of cells were prefixed with buffered glutaraldehyde (2.5% v/v) and 10 µl of prefixed cells were spotted onto poly-L-lysine coated glass slides (Corning BioCoat, USA). The glass slides were dried at room temperature and washed three times in 0.1 cacodylate and 5 g/l sucrose buffer. Washed slides were postfixed in a 1% (w/v) osmium solution for 40 min and rewashed three times with the cacodylate sucrose buffer. Subsequently, the slides were dehydrated in an ethanol series (30, 50, 70, 90, 96, and 100% ethanol in distilled water) for 5 min each, followed by two additional washing steps in 100% ethanol. After dehydration, the slides were dried with 100% ethanol using a Critical Point Dryer 300 instrument (Leica), mounted onto stubs, and gold sputter-coated (sputter coater JFC-2300HR, JOEL). Images were obtained with a scanning electron microscope (JSM-IT300, JOEL).

Paraformaldehyde fixation of LT0009 cells from an actively growing culture, fluorescence *in situ* hybridization (FISH) with 5'-end-mono-labeled rRNA-targeted probes (Biomers, Ulm, Germany) (Supplementary Table 1), and counter-staining with 4'-6-diamidino-2-phenylindole (DAPI) was performed as previously described [5]. A mouse colon tissue section was obtained from a previous study [6]. Cells and the tissue section were imaged using a confocal laser scanning microscope (Leica TCS SP8X, Germany) and FISH pictures were analyzed using the image analysis software daime [7]. A new probe TAU1151 was designed for LT0009 and related 16S rRNA sequences using the SILVA NR99, release 132 16S rRNA database [8], and probe design tools of the ARB program [9]. The hybridization buffer formamide concentration for optimal specificity and sensitivity of probe TAU1151 was determined by melting curve analysis (Supplementary Fig. 1). Probe MAIL1151 was designed as a competitor probe to eliminate nonspecific binding of TAU1151 to closely related *Mailhella* species. The SILVA database SSU_r138.1_REG was utilized to evaluate perfect-match coverage of the designed FISH probes using TestProbe 3.0 [10]. For increased specificity, MAIL1151 and TAU1151 should be used with different fluorophores and simultaneously in equimolar concentration.

**Genome sequencing and annotation**

Genomic DNA of strain LT0009 was extracted using the Wizard Genomic DNA purification Kit (Promega, USA) according to the manufacturer's procedures for both HiSeqV4 PE125 (Illumina) and MinION sequencing (Oxford Nanopore Technologies, Oxford, UK). Illumina and Nanopore sequences were demultiplexed, followed by adapter and barcode trimming using qCAT v. 1.1.0 (https://github.com/nanoporetech/qcat). A hybrid assembly of both Illumina and Nanopore reads was performed using Unicycler v. 0.4.6 [11]. The genome was annotated using the MicroScope annotation platform [12], and genes of interest were manually curated using the tools integrated into the MicroScope annotation platform (https://mage.genoscope.cns.fr/) as described previously [13]. Briefly, proteins annotated as homologous to proteins with a known function had an amino acid identity ≥40% (over ≥80% of sequence coverage) to a Swiss-Prot [14] protein or manually curated protein. Proteins annotated as putative homologs of the respective database entries had an amino acid identity ≥25% (over ≥80% of sequence coverage) to a Swiss-Prot or TrEMBL [15] entry. Hydrogenase genes were detected by the MicroScope annotation platform and further classified into different subgroups using the HydDB database tool [16].

**Phylogenetic and phylogenomic analyses**

The full-length 16S rRNA gene was retrieved from the genome of LT0009. Related 16S rRNA gene sequences with ≥80% similarity to LT0009 were recovered from the National Center for Biotechnology Information (NCBI) standard nucleotide database [17] and the SILVA database v.138 [10] using BLAST. 16S rRNA gene sequences of *Desulfovibrionaceae* type strains were extracted from List of Prokaryotic names with Standing in Nomenclature [18]. 16S rRNA gene sequences with ≥1,400 bp were dereplicated to remove redundant, 100% identical sequences, aligned using MUSCLE (v3.8.31) [19], and trimmed using TrimAl (v1.4. rev15) [20] with a gap threshold of 0.9. Maximum likelihood treeing of 16S rRNA gene sequences was performed using IQ-TREE (v. 1.6.2) [21] with 1,000x bootstrapping and model TVMe+R5. Sequence source environments were manually compiled from the NCBI SRA entries (Supplementary Table 2).

NCBI nucleic acid sequences with >60% coverage and >70% sequence similarity to *dsrAB* of LT0009 were recovered for phylogenetic analyses. Additional *dsrAB* sequences of related metagenome assembled genomes (MAGs) Mouse_MAG_UBA8003 (GCA_003512875.1) and *Mailhella massiliensis* strain Marseille-P3199 (GCA_900155525.1) were included in the phylogenetic analyses. Selected *dsrAB* nucleic acid sequences were aligned using MUSCLE (v3.8.31) [19] and trimmed using TrimAl (v1.4. rev15) [20] with -gt 0.1. The maximum likelihood tree was constructed using IQ-TREE (v1.6.2) [21] with 1000x bootstrapping [22] and automatic model selection.

Amino acid sequences related (>40% coverage, >70% sequence identity) to the *dsrE*-like gene (TAU_v1_1364) and rhodanese-like gene *sbdP* (TAU_v1_1430) were identified by BLAST and retrieved from NCBI for phylogenetic analyses. Amino acid sequences for *dsrEFH* genes (TAU_v1_1695, TAU_v1_1696, and TAU_v1_1697) were identified by hits to TIGRfam [23] and EggNOG [24] using HMM with e-value cutoff $1e^{-5}$. Additional DsrE, DsrEFH, and SbdP amino acid sequences were collected from the genomes in this study (Fig. 1b) using blastP with a minimum bit score of 100. CysH amino acid sequences were retrieved from NCBI using BLASTP (90% coverage and 65% sequence identity to CysH of *E. clostridioformis* YL32). Selected protein sequences were aligned with MAFFT (v7.475) [25], and the alignment was trimmed with TrimAl (v1.4.rev15) [20,25] with the flag -automated 1. Maximum-likelihood trees were created using the IQ-TREE web-server [26] with automatic substitution model selection and ultrafast bootstrapping (1000x) [22]. The trees were visualized with iTOL [27].

Complete genome sequences of representative *Desulfovibrionaceae* strains were retrieved from NCBI and representative high-quality *Desulfovibrionaceae* MAGs were selected from the integrated mouse gut metagenome catalog (iMGMC) [28] for phylogenomic analyses. Phylogenomic treeing with the IQ-TREE ML method (v. 1.6.2, model: LG+R3 as chosen by automatic model and 1000 ultrafast bootstrap runs [22]) was based on 43 phylogenetic marker protein sequences that were aligned and concatenated using CheckM. Average amino acid identity (AAI) and whole-genome average nucleotide identity (gANI) were calculated using the Enveomics Collection [29] and FastANI (v. 1.2) [30]. Genes involved in taurine (*tpa*), DHPS (*hpsGH*, *hpsO*, *hpsN*, *dphA*), 3-sulfolacetaldehyde (*slaB*), 3-sulfolactate (*suyAB*, *slsC*, *comC*), sulfoacetaldehyde (*xsc*, *sarD*), isethionate (*islAB*) and sulfite (*dsrABC*) metabolism were identified as described previously [31].

Presence of genes encoding thiosulfate reductases in LT0009 and *B. wadsworthia* genomes was revealed using blastP with an e-value cutoff of $1e^{-10}$ and minimal identity of 50%. Reference sequences included thiosulfate reductase PhsA from *Salmonella typhimurium* [32], thiosulfate reductase from *Nitratidesulfovibrio vulgaris* strain *Miyazaki F* and strain *Hildenborough* [33], Sox multienzyme system and thiosulfate dehydrogenase TsdA from *Paracoccus thiocyanatus* SST [34], Hdr-like enzyme from *Hyphomicrobium denitrificans* [35], rhodanese-like sulfurtransferase SbdP from *Aquifex aeolicus* [36], thiosulfohydrolase SoxB, and sulfur dioxygenase SdoAB homologs from *Erythrobacter flavus* 21-3 [37].

**Proteomics**

Harvested cells were disrupted using a UP50H – Compact Lab Homogenizer (Hielscher Ultrasound Technology, Germany) at cycle 0.5 and amplitude of 100%. Cell debris was removed by centrifugation (10 min, 10,000 x *g*) and the crude cell extracts were submitted to total proteomic analysis at the Proteomics Centre of the University of Konstanz as described previously [38]. The samples were analyzed on an Orbitrap Fusion with EASY-nLC 1200 (Thermo Fisher Scientific), and Tandem mass spectra were searched against the proteins inferred from the LT0009 genome using Mascot (Matrix Science) and Proteome Discoverer v 1.3 (Thermo Fisher Scientific) with Trypsin enzyme cleavage, static cysteine alkylation by chloroacetamide, and variable methionine oxidation.

**Transcriptomics**

RNA from triplicate LT0009 cultures grown with 10 mmol/l of taurine, sulfolactate, or thiosulfate was extracted using the Analytic Jena innuprep RNA Mini Kit 2.0 following manufacturer's instructions (JMF project JMF-2012-1). Extracted RNA was subjected to Turbo DNAse (Ambion) treatment to remove residual DNA contamination. Ribosomal RNA was not depleted to minimize sample processing biases. Stranded total RNA libraries were prepared using the NEBNext Ultra II Directional RNA Library Prep Kit for Illumina following the manufacturer's instructions. Paired-end (150 cycles) sequencing was performed on the HiSeq 3000 (Illumina). Raw reads were quality-filtered by removing adaptor-contaminated and low-quality reads at a Phred score of 28 using the bbduk function of BBMap (version 37.61) [39]. Next, filtered sequences were mapped as paired reads with a minimal identity of 99% to a reference file of all open reading frames of the LT0009 genome using the bbmap function of BBMap.

Normalized expression levels of transcripts were depicted as transcripts per million (TPM) values [40]. Read count values were used as input data for differential expression analysis by DESeq2 [41].

##### *Taurinivorans muris*- and *Bilophila wadsworthia*-related sequences in 16S rRNA gene amplicon datasets of human and animal guts

Occurrence and prevalence of 16S rRNA gene sequences related to LT0009 and *B. wadsworthia* ATCC 49260 were analyzed with the Integrated Microbial Next-Generation Sequencing (IMNGS) platform [42]. All Sequence Read Archive (SRA) amplicon sequence datasets containing the word "gut" in the "Origin" field were used for further analyses. A 97% sequence similarity cut-off was used for identification of related 16S rRNA gene sequences in the "gut" dataset that contained approximately 5.3 billion sequences from 123,723 gut samples, including 81,501 gut samples with host information. Further information on mouse studies with at least 20 samples that were positive for *B. wadsworthia* was manually compiled from the NCBI SRA entries or the corresponding publications (Supplementary Table 3).

##### Reanalysis of fecal 16S rRNA gene sequence data of wildR mice showing enhanced resistance against *Klebsiella pneumoniae*

We reanalyzed 16S rRNA gene amplicon sequencing data of the 2nd (PRJNA390686) [43] and the 10th generation (PRJNA666931) [44] of wildR mice (a specific-pathogen-free mouse colony whose germ-free founders received the microbiota of wild mice) that showed increased representation of *Deltaproteobacteria* and colonization resistance against the enteropathogen *Klebsiella pneumoniae* in a previous study [44]. Downloaded fastq files were run through the DADA2 pipeline [45]. Briefly, the sequences were filtered with parameters maxN=0, maxEE=c (2,2), truncQ=10, and truncLen=c (180, 150). Following chimera removal, taxonomy was assigned to the amplicon sequence variants (ASVs) using the silva_nr99_v138 reference database [10]. ASVs assigned to unclassified *Desulfovibrionaceae* spp. were blasted against the 16S rRNA gene sequence of strain LT0009. ASVs with a similarity of >98% to the LT0009 16S rRNA gene sequence were re-classified as *T. muris*.

##### Gnotobiotic mouse experiments

Twelve germ-free C57BL/6 mice were obtained from Dr. Basic (Hannover Medical School, Hannover, Germany) for the colonization experiment. The animal experiment was approved by the local authorities (Regierung von Oberbayern; ROB-55.2-2532.Vet_02-20-84). The synthetic, 12-member Oligo-Mouse-Microbiota ($OMM^{12}$) community consists of *Acutalibacter muris* KB18, *Akkermansia muciniphila* YL44, *Bacteroides caecimuris* I48, *Bifidobacterium* animalis YL2, *Blautia coccoides* YL58,

*Enterocloster clostridioformis* YL32, *Clostridium innocuum* I46, *Enterococcus faecalis* KB1, *Flavonifractor plautii* YL31, *Limosilactobacillus reuteri* I49, *Muribaculum intestinale* YL27, and *Turicimonas muris* YL45. Strain LT0009 was cultivated in Anaerobic Akkermansia Medium [46] supplemented with 10 mmol/l taurine, 20 mmol/l sodium pyruvate, and 200 µg/l naphthoquinone. A subculture was incubated at 37°C for 3 days. Mice (n=6) colonized with OMM[12] strains were orally (50 µl) and rectally (100 µl) inoculated with the LT0009 subculture and the control group (n=6) was treated with the same volume of sterile 1x phosphate-buffered saline. After 10 days, the mice were infected with the human enteric pathogen *Salmonella enterica* serovar Typhimurium (avirulent *S. enterica* Tm strain M2702; $5\times10^7$ c.f.u.). At the same time, the fecal microbiota composition was determined by strain-specific qPCR assays as previously described [47]. New 16S rRNA gene-targeted primers (forward: 5'-TTCGGATCGTAAACCTCTGTCA-3'; reverse: 5'-GGTACCGTCAATTCACTGCTGAT-3') and a detector probe (5' 6-carboxyhexafluorescein-CAGGGAAGAACGGTCAC-black hole quencher 1-3') for qPCR of LT0009 were designed using Primer Express 3 (Applied Biosystems, Life Technologies). The mice were sacrificed by cervical dislocation two days post infection (p. i.). Abundance of viable *S. enterica* Tm at 24 and 48 h p.i. in the feces and at 48 h p.i. in the cecal content was determined by plating [47].

Fecal samples of three mice from each group on day two p.i. were selected for metatranscriptome sequencing (JMF project JMF-2104-01). RNA was extracted using the Analytic Jena innuprep RNA Mini Kit 2.0 following manufacturer's instructions. Extracted RNA was subjected to Turbo DNAse (Ambion) treatment to remove residual DNA contamination. Ribosomal RNA was depleted using the RiboZero rRNA depletion Kit (Illumina), and stranded RNA libraries were prepared using the NEBNext Ultra II Directional RNA Library Prep Kit for Illumina following manufacturers' instructions, and sequenced on an Illumina Novaseq 6000 in paired-end mode (2 x 100 bp). Reads were filtered for contamination and adapters using BBDuk as follows: k=23, mink=11 (https://sourceforge.net/projects/bbmap/). Reads after quality control were mapped to the reference genomes of individual strains [48] and *S. enterica* Tm (GCA_000210855.2) (Supplementary Table 4) using BBMap (version 38.92) [39] at 98% sequence identity. Genes of OMM[12] strains and *S. enterica* Tm that were significantly differentially expressed between mice with or without strain LT0009 were revealed using Deseq2 [41]. Sequences of eight representative bile salt hydrolase (BSH) genes from the human gut microbiome [49] were used to produce the BLASTP search database and used to identify BSH homologs with >30% identity in genomes of the gnotobiotic community (Supplementary Table 5). Prophage regions in *E. clostridioformis* YL32 were predicted using PHASTER [50].

**LT0009-centric gut metatranscriptome analyses of laboratory mice**

Cecal and fecal metatranscriptomes from a high-glucose diet experiment in mice (HG study, Hanson et al., unpublished) (JMP project JMF-2101-5) were analyzed for LT0009 gene expression. Mouse experiments were conducted following protocols approved by Austrian law (BMWF-66.006/ 0032-WF/V/3b/2014). Six seven-week-old C57BL/6 mice were randomly allocated to groups and fed either a control diet (5% glucose; n=3) or a high-glucose diet (65% glucose; n=3) for about 3 weeks. Cecal and fecal contents were collected for RNA extraction. RNA was extracted as previously described [51]. Ribosomal RNA was depleted using the RiboZero rRNA depletion Kit (Illumina), and stranded RNA libraries were prepared using the NEBNext Ultra II Directional RNA Library Prep Kit for Illumina following manufacturers' instructions, and sequenced on an Illumina HiSeq 3000 in paired-end mode (75 + 91 bp). Library preparation, sequencing, and data processing were performed at the Joint Microbiome Facility of the Medical University of Vienna and the University of Vienna.

We additionally re-analyzed mouse gut metatranscriptomes from a previous study for LT0009 gene expression (Plin2 study) [52]. Sequence data (PRJNA379425) derived from eight-week-old C57BL/6 wild-type and Perilipin2-null (Plin2) mice fed high-fat/low-carbohydrate or low-fat/high-carbohydrate diets. Low-quality reads were removed at a Phred score of 28 using the bbduk function of BBMap (version 37.61) and filtered sequences were mapped to the LT0009 genome using the bbmap function of BBMap with a 99% similarity cutoff [39]. Expression levels of transcripts were normalized as TPM values for comparison.

#### Results & discussion

##### Additional energy metabolism of *Taurinivorans muris* LT0009

Genome reconstruction of LT0009 suggested the potential to utilize lactate, pyruvate, and $H_2$. We experimentally confirmed that lactate and pyruvate are used as electron donors for taurine respiration (Fig. 2b). LT0009 expressed lactate permease LutP and L-lactate dehydrogenase LutABC/LldEFG for lactate import and oxidation to pyruvate (Fig. 2a, Supplementary Table 7) [53,54]. A homolog of another putative lactate dehydrogenase, the flavin and iron–sulfur containing membrane-associated oxidoreductase Dld-II [55] is also encoded by LT0009 (TAU_v1_1652), yet was not expressed (Supplementary Table 7). Pyruvate can be further oxidized by LT0009 to acetyl-CoA with either pyruvate:ferredoxin oxidoreductase Por or pyruvate formate-lyase PflD. Growth experiments of LT0009 with or without pyruvate revealed that additional pyruvate can stimulate its growth (Supplementary Fig. 3b).

For comparison, *B. wadsworthia* strains can grow with lactate, pyruvate, formate, and $H_2$ as electron donors [38,56,57]. Strain LT0009 also used formate as electron donor for taurine-respiration (Fig. 2b), yet its formate metabolism remains unresolved as genes for formate dehydrogenase or the formate hydrogen-lyase complex were not detected. LT0009 encodes four hydrogenases (Supplementary Table 6). The presence of genes for a respiratory, $H_2$-uptake group 1b [NiFe]-hydrogenase (*hybAC*) and its corresponding maturation factor *hypABCDEF* is consistent with the prevalence of this enzyme in hydrogenotrophic *Desulfobacterota* in the gut [58] and suggested $H_2$ as an additional electron donor for LT0009. However, strain LT0009 did not grow with $H_2$ as electron donor under the conditions we used (Fig. 2b). Besides the potential for $H_2$-utilization, LT0009 also encodes a fermentative group A1 [FeFe] hydrogenase and a group C3 [FeFe] hydrogenase of yet undetermined biochemical function [16]. The presence of *cooMKLXUHF* operon encoding $H_2$-evolving group 4c [NiFe] carbon monoxide-induced hydrogenase suggested carbon monoxide, a ubiquitous molecule in the gut [59], as a potential electron donor and source of $H_2$ for LT0009. However, LT0009 lacks the gene cluster *cooFSCTJ* that encodes carbon monoxide dehydrogenase CooS and the electron-transfer protein CooF, which are required for optimal $H_2$ production by the carbon monoxide-induced hydrogenase in *Rhodospirillum rubrum* [60,61].

LT0009 encodes a complete glycolysis/gluconeogenesis pathway. The coupling of electron transfer to energy conservation is likely mediated by an $H^+$/$Na^+$-pumping Rnf complex (RnfCDGEAB) [62] and ATP synthase (AtpABCDEFGH).

**Description of *Taurinivorans* gen. nov.**

*Taurinivorans* gen. nov. (Tau.ri.ni.vo'rans. N.L. n. *taurinum*, taurine; L. part. adj. *vorans*, eating; N.L. masc. n. *Taurinivorans*, a taurine eater). Comparative genome analyses suggest the common electron acceptor is taurine, which is degraded and reduced to sulfide via the Tpa-Xsc-DsrAB-DsrC pathway. Type species: *Taurinivorans muris* sp. nov., family: *Desulfovibrionaceae* VP, order: *Desulfovibrionales* VP (T) emend., class: *Desulfovibrionia* class. nov., phylum: *Desulfobacterota* phyl. nov. [63].

**Description of *Taurinivorans muris* sp. nov.**

*Taurinivorans muris* sp. nov. (mu'ris. L. gen. n. *muris*, of a mouse, referring to its origin from the mouse intestine). The type strain is strain LT0009$^T$ (=DSM 111569 =JCM 34262), isolated from the mouse gut with taurine as the electron acceptor and lactate/pyruvate as electron donors. Formate was also used as an electron donor for taurine respiration. Cells are Gram-stain-negative, spirilloid in shape, and motile by means of lophotrichous polar flagella. The temperature range is 27-42°C and the optimum pH is 6.5 (range 6-8.5) for strictly anaerobic growth. The optimal taurine concentration for growth is

40 mmol/l, higher taurine concentrations inhibited growth. Sulfolactate and thiosulfate are additional electron acceptors for anaerobic respiration and are also reduced to hydrogen sulfide. Yeast extract and 1,4-naphthoquinone are required as growth supplements for laboratory cultivation of the isolate. Its genome size is 2.2 Mbp with a G+C content of 43.6%. The GenBank accession numbers for the genome and the 16S rRNA gene sequence of strain LT0009$^{T}$ are CP065938 and MW258658, respectively.

#### Supplementary table legends

**Supplementary Table S1.** 16S rRNA-targeted oligonucleotide probes used for FISH analysis in this study.

**Supplementary Table S2.** Accession numbers and source information of the sequences used for 16S rRNA phylogenetic analysis, as collected from NCBI (https://www.ncbi.nlm.nih.gov).

**Supplementary Table S3.** Additional information for *B. wadsworthia*-positive samples from the mouse gut.

**Supplementary Table S4.** Gnotobiotic mouse microbiota genomes.

**Supplementary Table S5.** Differential expression of bile salt hydrolase genes in the $OMM^{12}$ community.

**Supplementary Table S6.** Curated annotation of selected LT0009 genes, with a focus on sulfur and energy metabolism. COG class IDs were assigned by MaGe (Cognitor, www.ncbi.nlm.nih.gov/COG/) and NOG IDs were assigned by the best-match principle [64,65].

**Supplementary Table S7.** Transcriptome and proteome analyses of *T. muris* LT0009. Columns show the locus tag, length of the gene in base pairs [bp], transcripts per million [TPM], log fold change, normalized protein expression areas and the annotation. TPM is given as an average of triplicate cultures grown with taurine (T), sulfolactate (SL), and thiosulfate (Thi). In addition, the p-values for the transcripts and protein expressions are provided, with *p*-values <0.01 highlighted in blue and *p*-values <0.05 highlighted in orange.

**Supplementary Table S8.** Prevalence of *Taurinivorans muris* and *Bilophila wadsworthia*-related sequences with 97% identity cut-off across 16S rRNA gene amplicon datasets of diverse hosts. Numbers in parentheses indicate the number of amplicon samples analyzed per host.

**Supplementary Table S9.** Gene expression of $OMM^{12}$ strains and *S. enterica* Tm SL1344 in gnotobiotic mice with and without LT0009. Gene annotations are derived from the reference genomes at NCBI. log2FC: logarithm to the base2 (Fold Change), padj: adjusted p-value obtained by DESeq2.

### Supplementary Figure S1

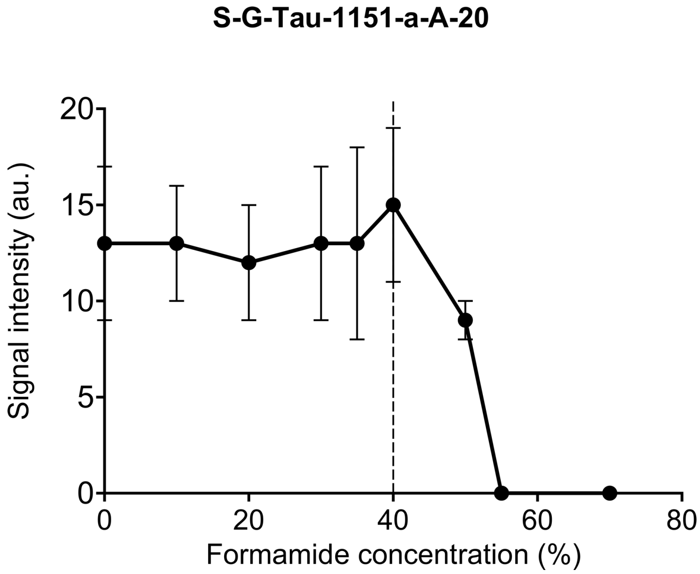


**Supplementary Figure 1. Formamide dissociation profile of FISH probe TAU1151.** Fluorescence signal intensities under increasing formamide concentrations are depicted for strain LT0009 hybridized with the Cy3-labeled probe TAU1151 for the genus *Taurinivorans*. The dashed vertical line indicates the formamide concentration for best possible specificity and sensitivity of probe TAU1151.

### Supplementary Figure S2

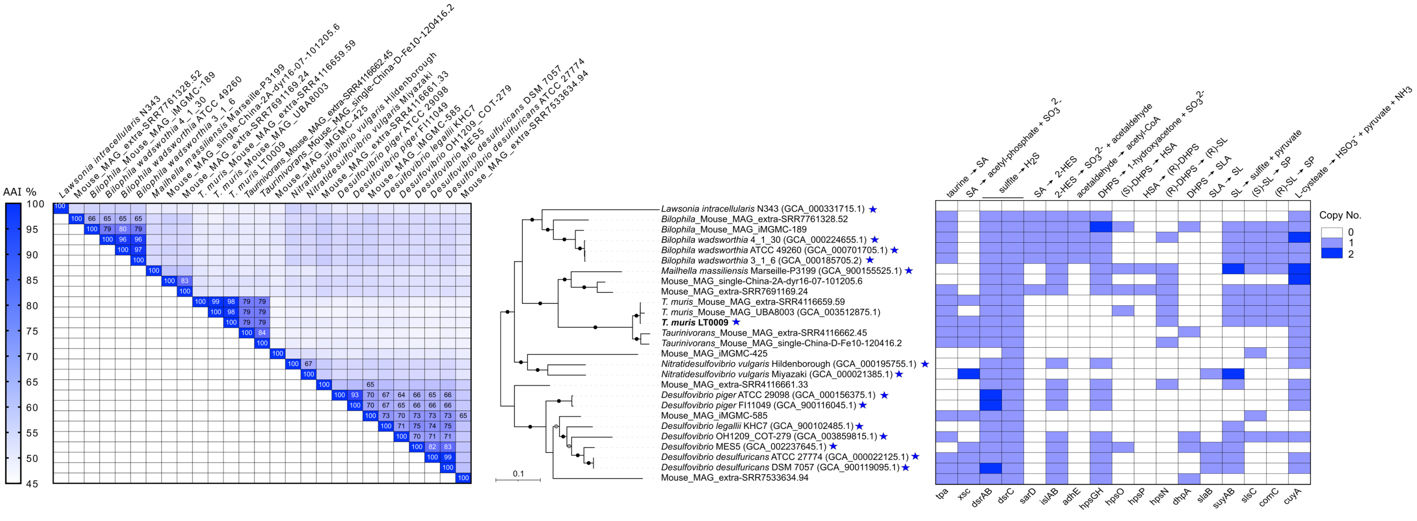


**Supplementary Figure 2. Average amino acid identities, phylogenomic relationship, and organosulfonate metabolism gene distribution of strain LT0009 and related genomes/MAGs of the *Desulfovibrio-Mailhella-Taurinivorans-Bilophila* lineage**. Left panel. Matrix of pairwise AAIs; only values ≥63.4 (genus cut-off [66]) are shown. Middle panel. Phylogenomic tree from Figure 1b is used here to illustrate the AAI and sulfur metabolism gene similarity of the close relatives. Right panel. Presence/absence of organosulfonates metabolism genes in the genomes of LT0009 and its relatives. Organosulfonate metabolism genes and encoded enzymes/proteins: *tpa*, taurine:pyruvate aminotransferase; *xsc*, sulfoacetaldehyde acetyltransferase; *dsrAB*, dissimilatory sulfite reductase subunits A and B; *dsrC*, sulfite reduction co-substrate DsrC; *sarD*, sulfoacetaldehyde reductase; *islAB*, isethionate sulfite-lyase complex; *ahdE*, CoA-acylating aldehyde dehydrogenase; *hpsGH*, DHPS sulfite-lyase complex; *hspNOP*, DHPS dehydrogenases; *dhpA*, $NAD^+$-dependent DHPS dehydrogenase; *slaB*, 3-sulfolactaldehyde dehydrogenase; *suyAB*, (R)-sulfolactate sulfo-lyase; slsC, (S)-sulfolactate dehydrogenase; comC, (2R)-3-sulfolactate dehydrogenase; *cuyA*, L-cysteate sulfo-lyase. SA, sulfoacetaldehyde; 2-HES, 2-hydroxyethane-1-sulfonic acid (isethionate); DHPS, 2,3-dihydroxypropane-1-sulfonate; HSA, 2-oxo-3-hydroxy-propane-1-sulfonate; SL, 3-sulfolactate; SLA, 3-sulfolactaldehyde; SP, sulfopyruvate.

### Supplementary Figure S3

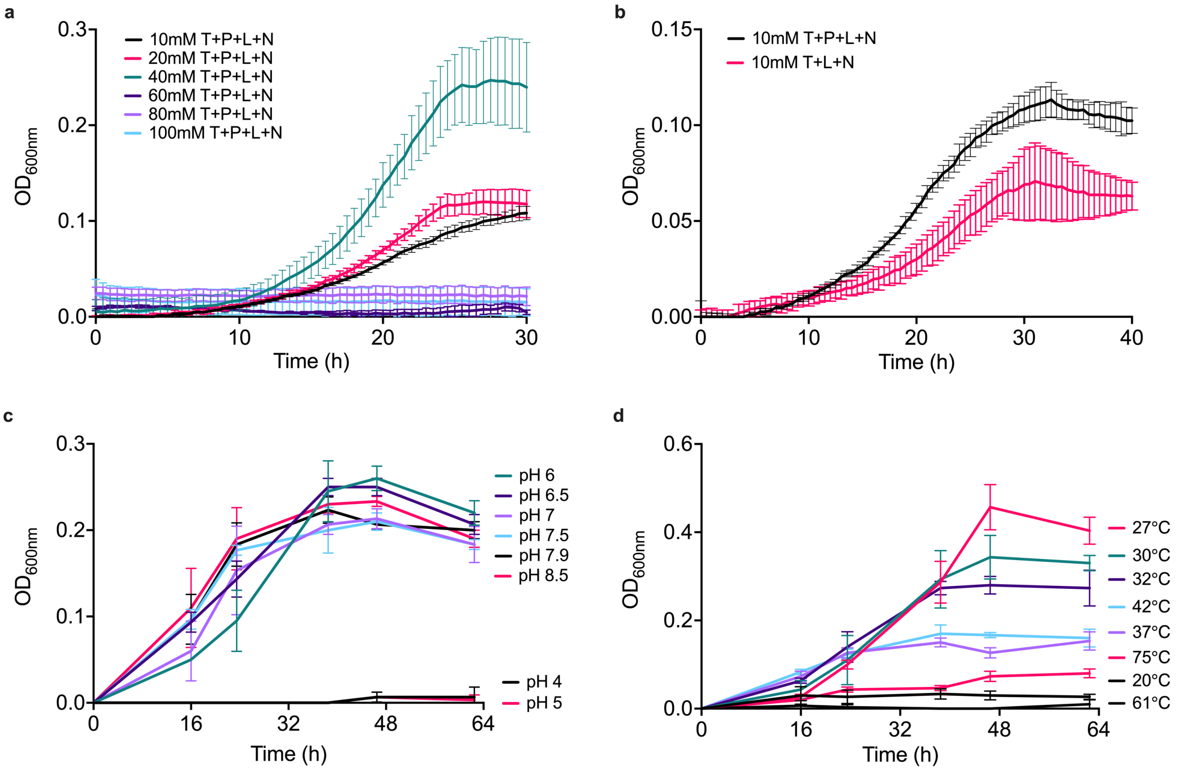


**Supplementary Figure 3. Anaerobic growth tests of *Taurinivorans muris* LT0009.** LT0009 grown **a.** with different taurine concentrations (10, 20, 40, 60, 80, and 100 mmol/l) and 20 mmol/l pyruvate, 10 mmol/l lactate, and 0.2 mg/l 1,4-naphthoquinone **b.** with or without pyruvate **c.** at different pH values (temperature 37°C) and **d.** at different temperatures (pH 7.2). Growth curves show the averages of optical density measurements at 600 nm ($OD_{600nm}$) in triplicate culture. Error bars represent one standard deviation. OD:. T, taurine; P, pyruvate; L, lactate; N, 1,4-naphthoquinone.

### Supplementary Figure S4

a

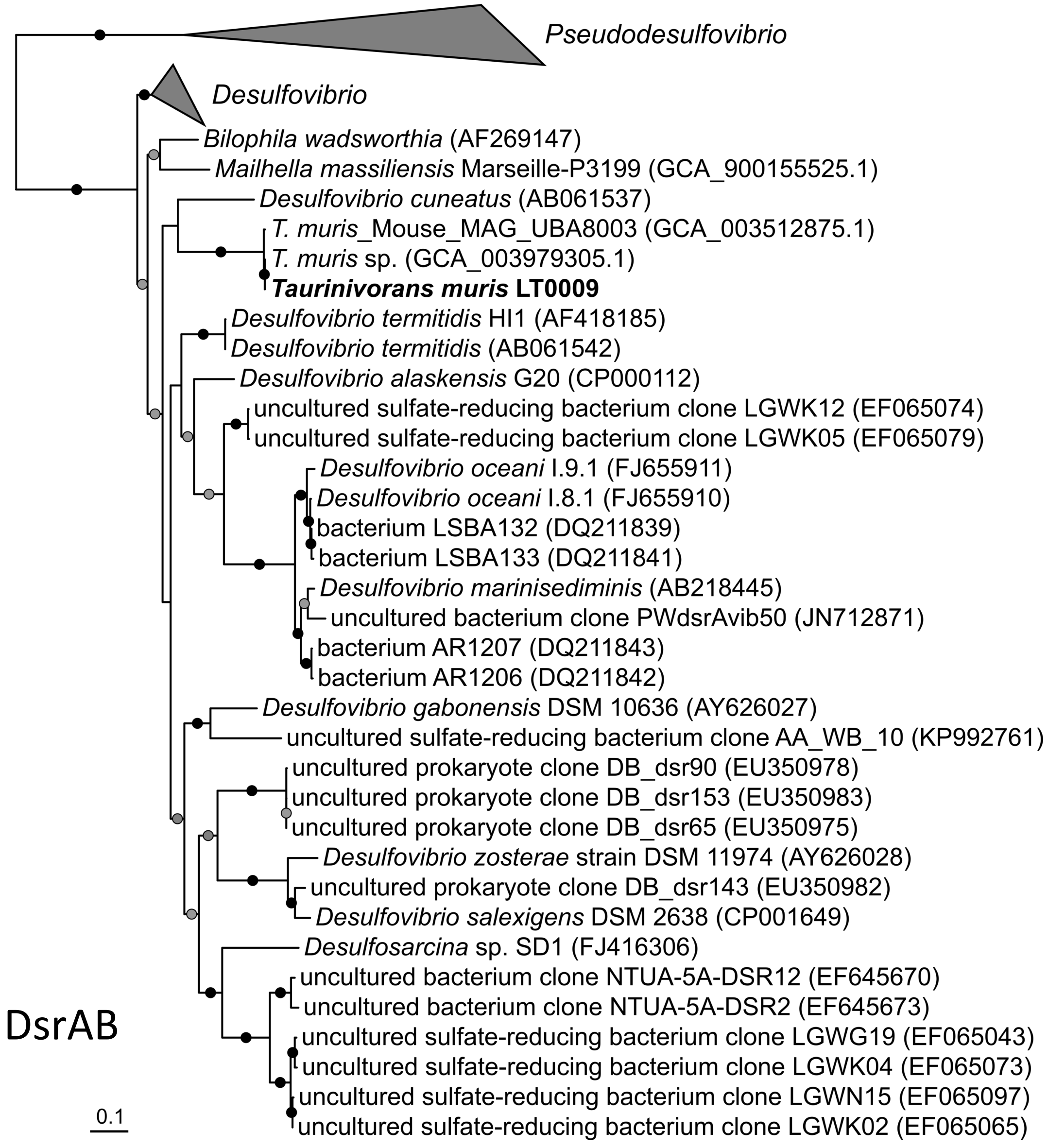

### Supplementary Figure S4

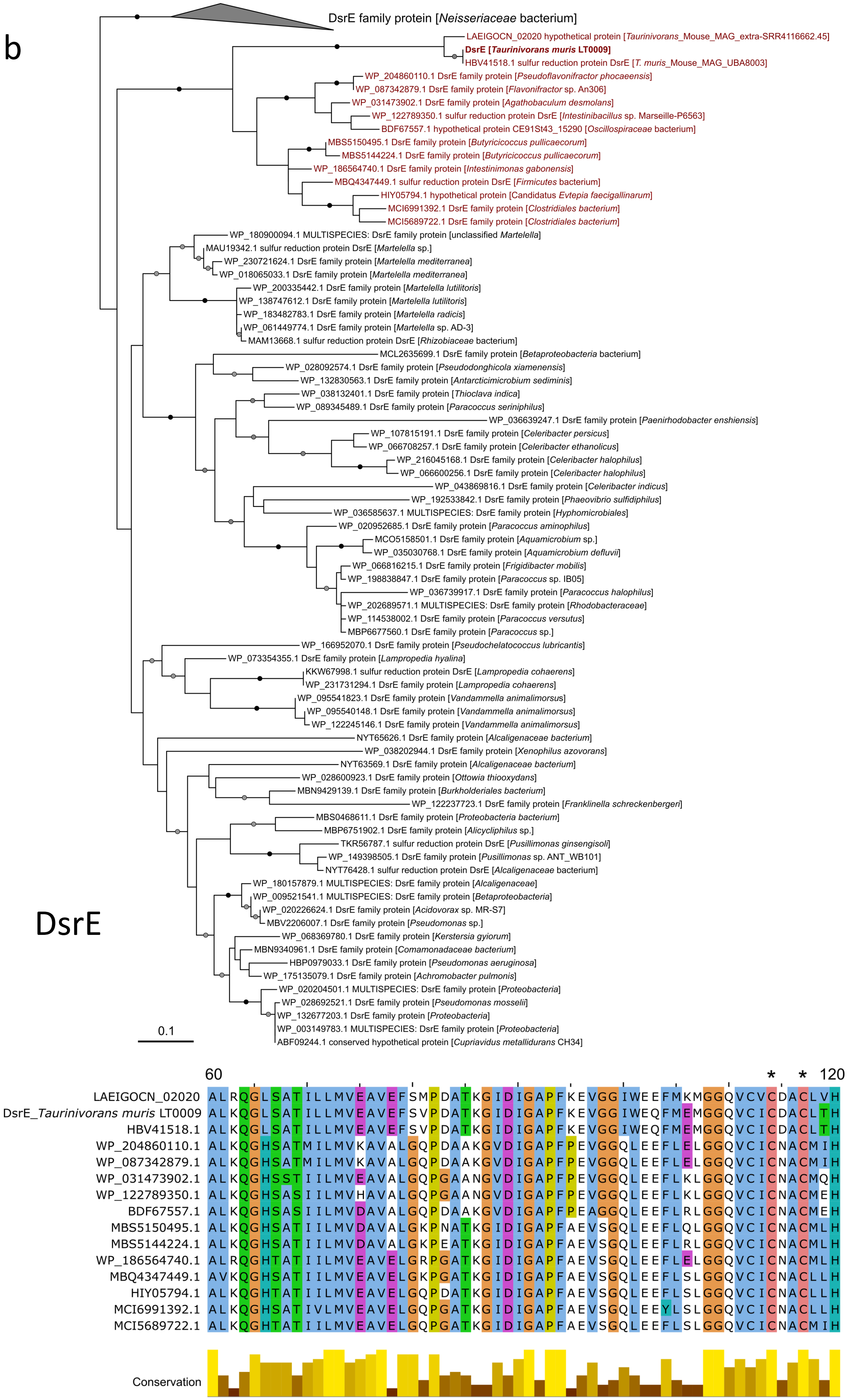

### Supplementary Figure S4

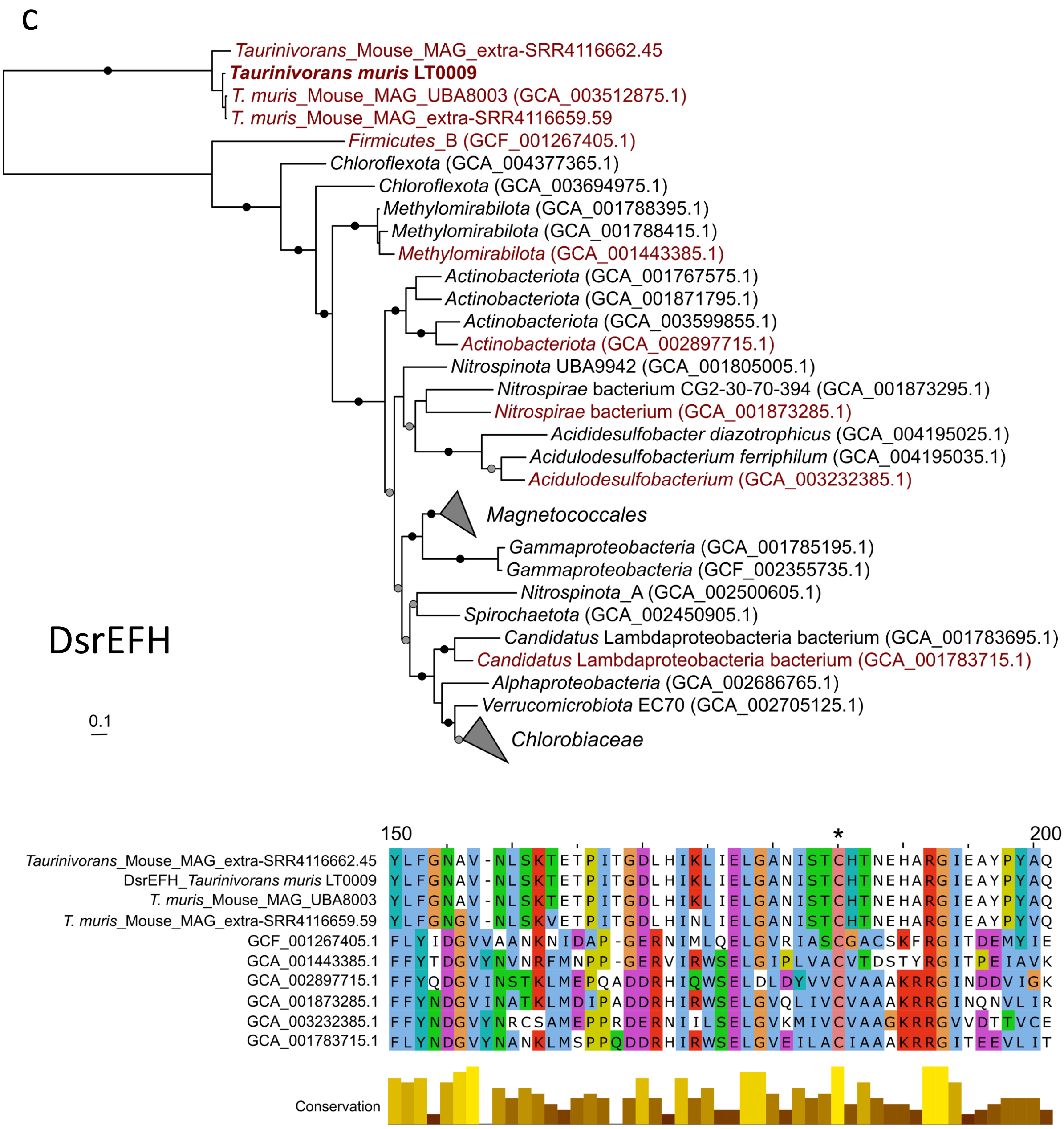

### Supplementary Figure S4

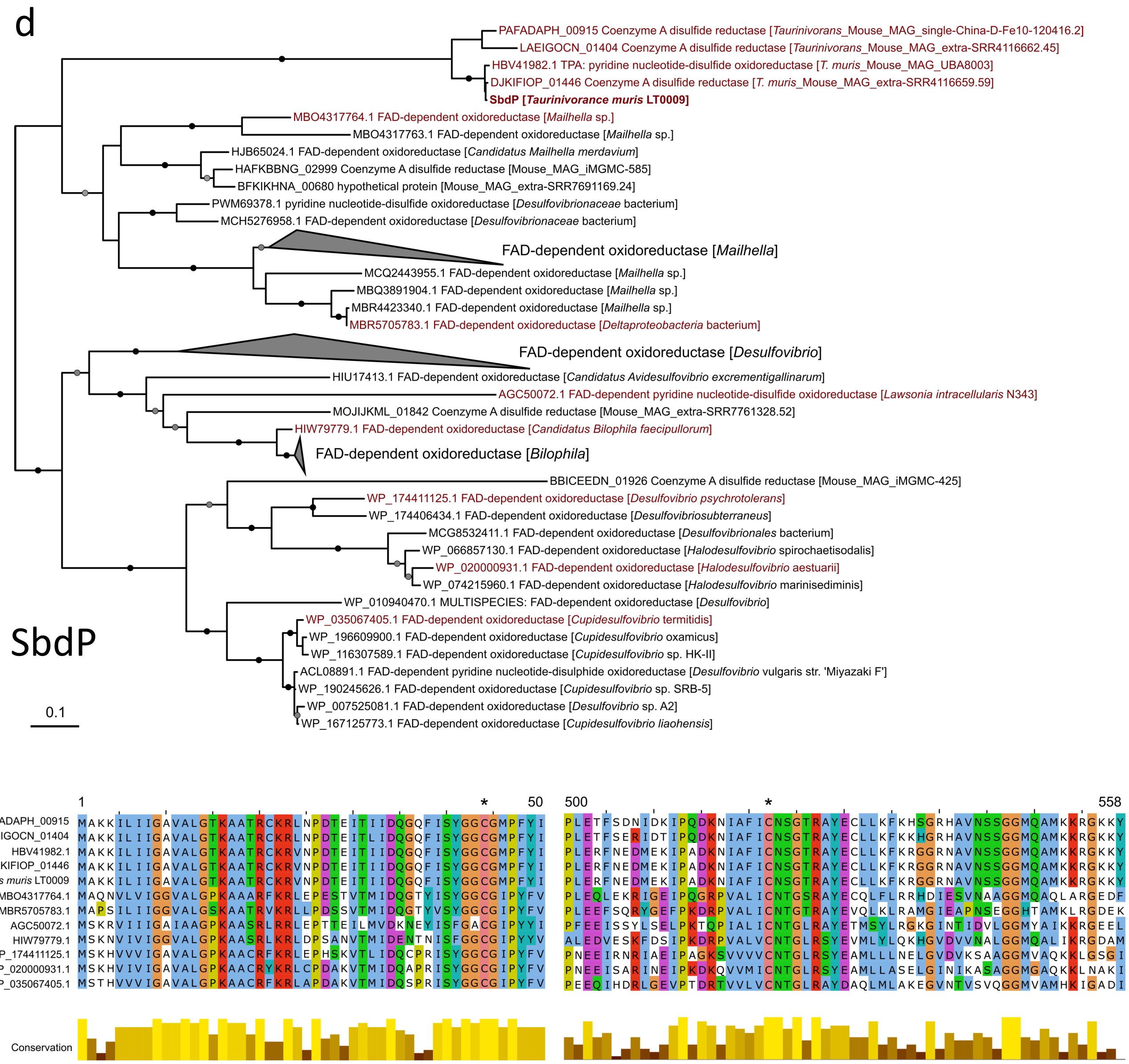


**Supplementary Figure 4. Phylogeny and conserved amino acid residues of selected sulfur metabolism proteins of *Taurinivorans muris* LT0009. a.** *dsrAB* nucleic acid tree. **b.** DsrE (TAU_v1_1364) protein tree. **c.** Concatenated DsrEFH protein tree. **d.** SdbP protein tree. *Taurinivorans muris* LT0009 is highlighted in bold and the scale bar represents 0.1 estimated substitutions per residue in all trees. All trees are midpoint rooted and bootstrap branch supports equal to or greater than 95% and 80% are indicated by black and grey circles, respectively. Amino acid alignments show select regions with active site cysteines (labeled with an asterisk) conserved across all sequences in the respective tree. The alignment is only shown for sequences marked in red in the tree. The alignment conservation profile is based on all sequences in the tree. The inferred DsrE-like amino acid sequence in *T. muris* LT0009 and related sequences contain a Cys-X2-Cys structure that is different from those in known thiosulfate-transferring DsrE homologs [67]. The cysteine site in DsrEFH is responsible for sulfur atom transfer from DsrEFH to DsrC in sulfur oxidizers [68]. The cysteine in SbdP is also involved in sulfur atom binding [36].

### Supplementary Figure S5

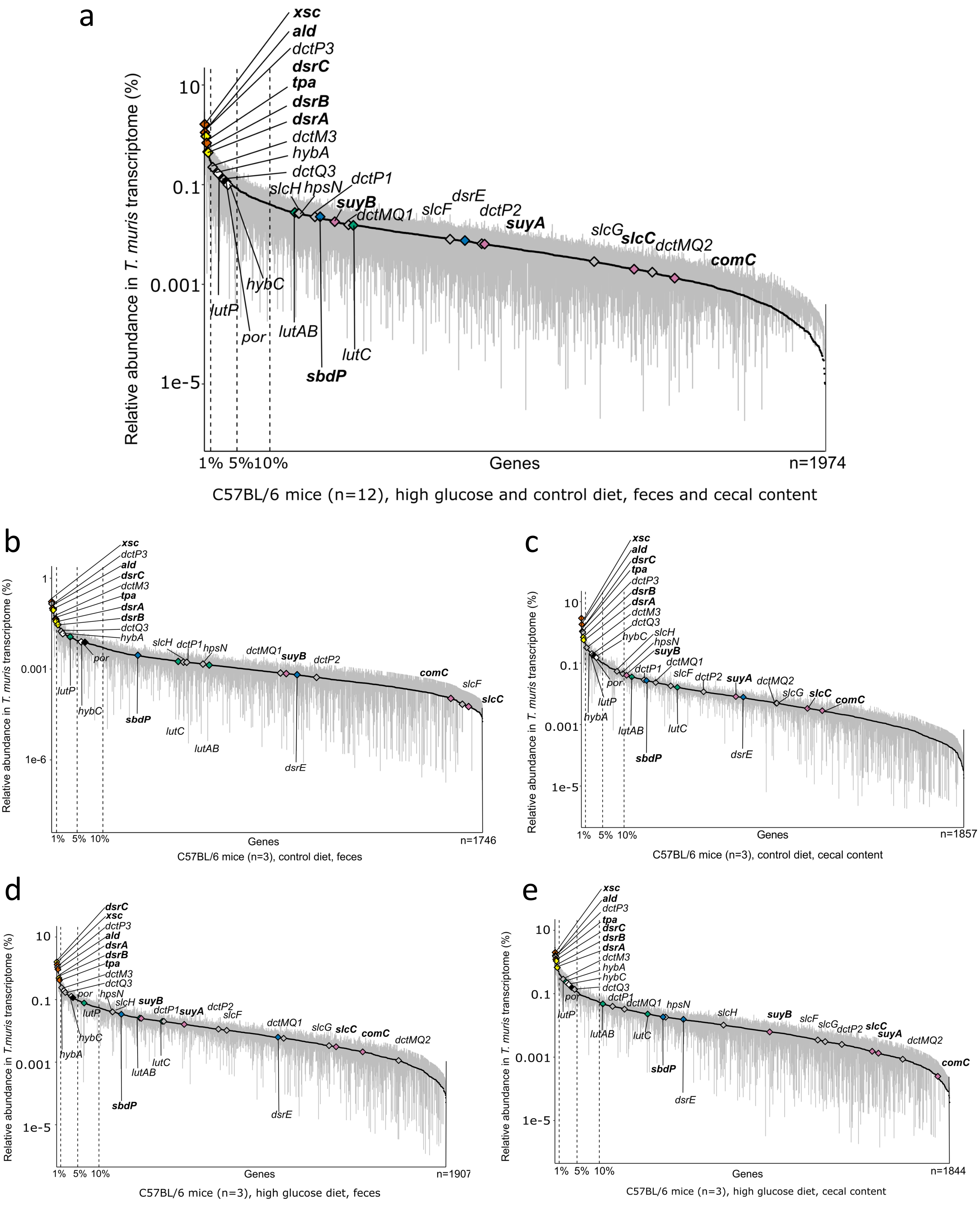

### Supplementary Figure S5

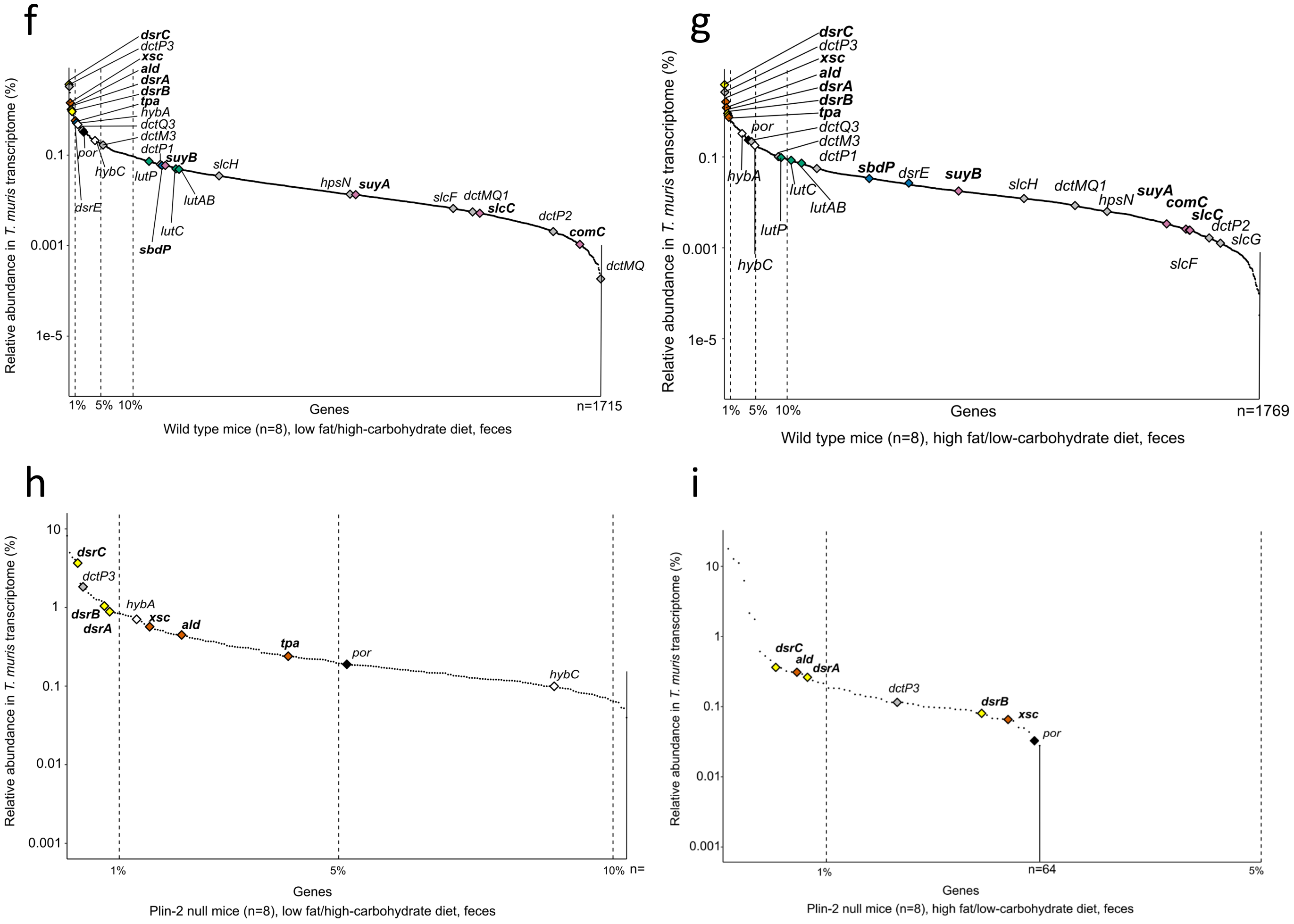


**Supplementary Figure 5. Mouse gut metatranscriptomics shows taurine respiration as the main *in vivo* metabolic niche of *Taurinivorans muris* independent of intestinal location, host genotype, and host diet.** LT0009-centric gut metatranscriptome analyses of laboratory mice from two studies: (**a**-**e**) HG study (Hanson et al. unpublished), (**f**-**i**) Plin2 study [52]. All plots show the ranked relative transcript abundance of LT0009 genes and the total number of transcribed LT0009 genes detected in the respective datasets. Genes for taurine (*tpa*, *xsc*, *ald*), sulfite (*dsrAB, dsrC*), sulfolactate (*suyAB*, *sclC*, *comC*), thiosulfate (*sbdP, dsrE*), pyruvate (*por*), lactate (*lutAB, lutC, luP*), and hydrogen (*hybA*, *hybC*) metabolism are shown in different colors. Sulfur metabolism genes are further highlighted in bold font. Vertical dashed lines indicate the top 1%, 5%, and 10% of the protein-coding genes in LT0009 genome (n=2059). For the HG study, each point in the rank abundance plot is the mean relative abundance of a gene transcript and error bars correspond to the 95% confidence interval of the mean. The metatranscriptome data from Plin2 study include data from eight animals in one SRA file; data for each individual mouse was not available. Thus, each point in the rank abundance plot represents the relative gene expression across all eight samples. **a.** Gene expressions in all mice from HG study (n=12). **b.** Gene expressions in fecal samples from mice fed a control diet (n=3). **c.** Gene expression in cecal samples from mice fed a control diet (n=3). **d.** Gene expressions in fecal samples from mice fed a high-glucose diet (n=3). **e.** Gene expressions in cecal samples from mice fed a high-glucose diet (n=3). **f.** Gene expression in fecal samples from wild type mice fed a low-fat/high-carbohydrate diet (n=8). **g.** Gene expression in fecal samples from wild type mice fed a high-fat/low-carbohydrate diet (n=8). **h.** Gene expression in fecal samples from Plin-2 null mice fed a low-fat/high-carbohydrate diet (n=8). **i.** Gene expression in fecal samples from Plin-2 null mice fed a high-fat/low-carbohydrate diet (n=8).

### Supplementary Figure S6

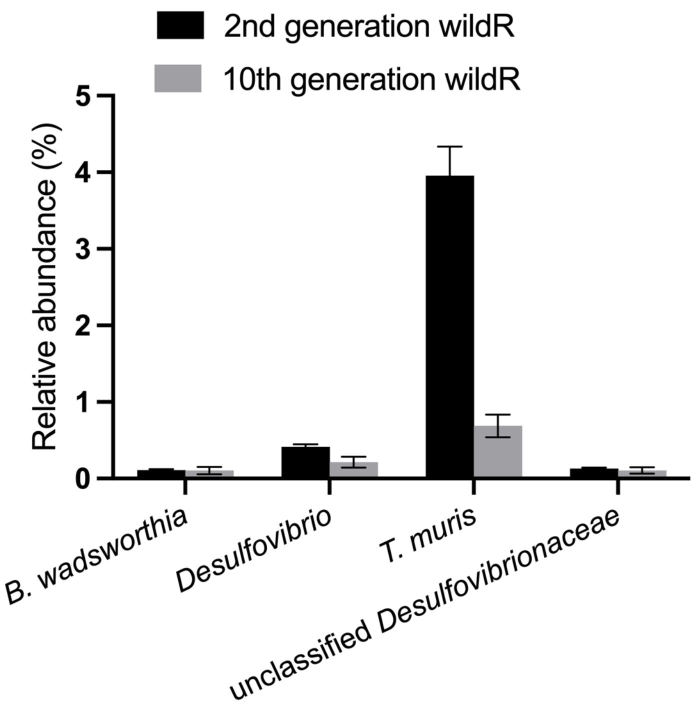


**Supplementary Figure 6. *Taurinivorans muris* is the dominant *Desulfovibrionaceae* member in wildR mice that provided enhanced, $H_2S$-mediated protection against *Klebsiella pneumoniae*.** A previous study showed that the 2nd and 10th generation of wildR mice (specific-pathogen-free mice whose germ-free founders received the microbiota of wild mice) displayed enhanced resistance against the enteropathogen *Klebsiella pneumoniae* compared to control specific-pathogen-free mice [44]. WildR mice showed increased abundance of *Proteobacteria*, particularly taurine-utilizing *Deltaproteobacteria* encoding the Tpa-Xsc-Dsr taurine metabolism. Enhanced colonization resistance was mediated by taurine-driven production of $H_2S$ that inhibited enzymes of pathogens for aerobic respiration and thus restricted their access to strictly respiratory substrates. Here, our reanalysis of the 16S rRNA gene amplicon sequencing data revealed that *T. muris* is the most abundant deltaproteobacterium in wildR mice with relative abundances of 4.0% and 0.7% of the total bacterial community in the 2nd and 10th generation, respectively.

### Supplementary Figure S7

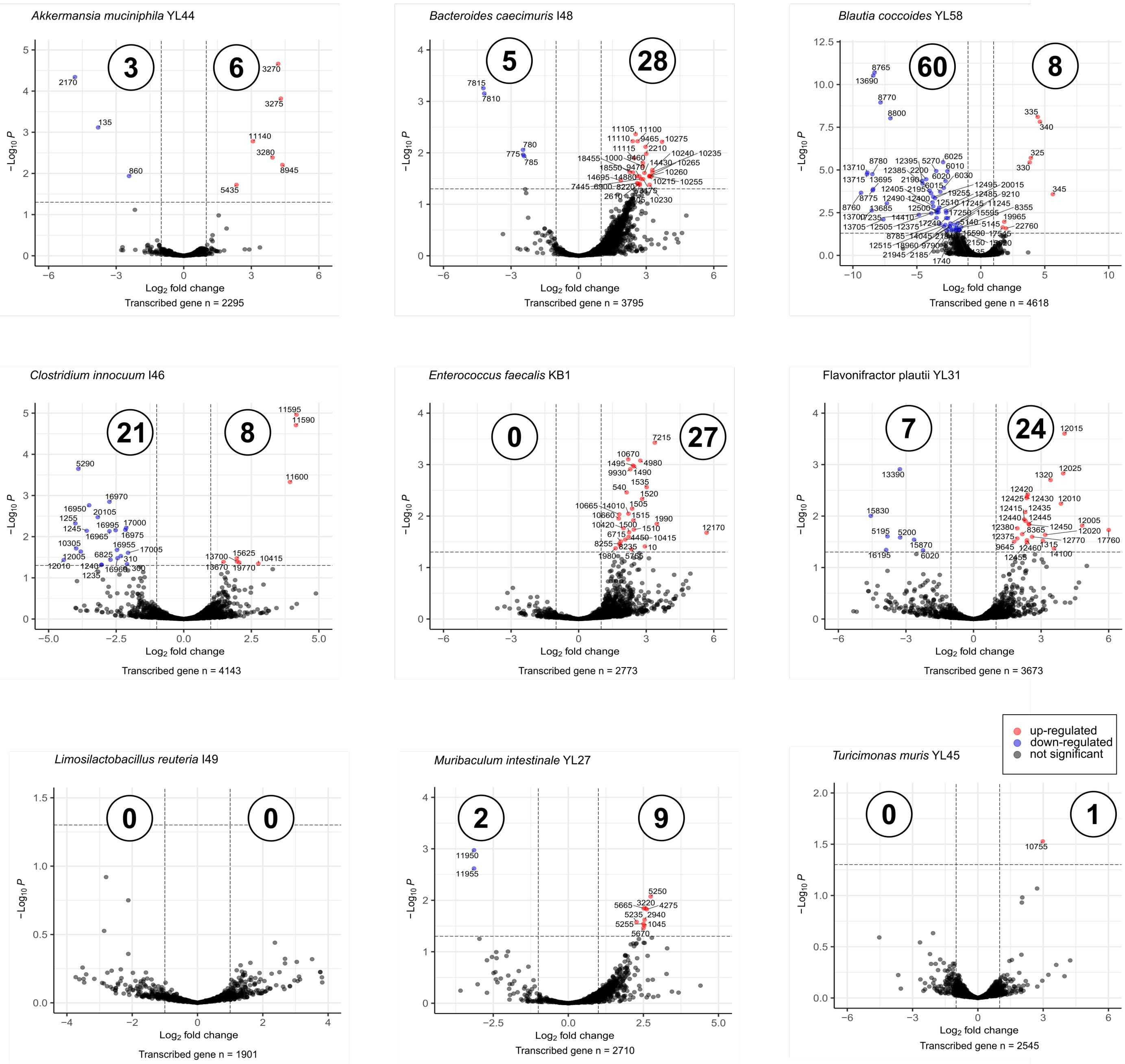


**Supplementary Figure 7. Differential gene expression of OMM$^{12}$ strains in gnotobiotic mice with and without *Taurinivorans muris* LT0009 and infected with *Salmonella enterica* Tm$^{avir}$ M2702.** No reads with 98% sequence identity to genomes of *A. muris* KB18 and *B. animalis* YL2 were detected, which is consistent with lack of evidence of colonization of these strains by qPCR (Fig. 4b). Volcano plots show differential gene transcription of individual OMM$^{12}$ strains in mice with and without LT0009. The x-axis shows log-fold-change in transcription and the y-axis the negative logarithm10-transformed adjusted p values. Red and blue dots show significantly (adjusted-p value <0.05) up-regulated (log2 fold change >1) and down-regulated (log2 fold change <-1) genes in mice with LT0009, respectively, and are labeled with locus tag numbers. Numbers in circles show the total numbers of up- and down-regulated genes per strain.
